# Perceived biodiversity: is what we measure also what we see and hear?

**DOI:** 10.1101/2024.04.03.587905

**Authors:** Kevin Rozario, Taylor Shaw, Melissa Marselle, Rachel Rui Ying Oh, Erich Schröger, Mateo Giraldo Botero, Julian Frey, Valentin Ștefan, Sandra Müller, Michael Scherer-Lorenzen, Bogdan Jaroszewicz, Kris Verheyen, Aletta Bonn

## Abstract

1. Biodiversity is crucial for human health and well-being. Perceived biodiversity - people’s subjective experience of biodiversity - seems to be particularly relevant for mental well-being.

2. Using photographs and audio recordings of forests that varied in levels of species richness, we conducted two sorting studies to assess how people perceive visual and acoustic diversity and whether their perceptions align with measured tree and bird species richness (‘actual diversity’). Per study, 48 participants were asked to sort the stimuli according to any similarity-based sorting criteria they liked (‘open sorts’) and perceived diversity (‘closed sorts’).

3. The main perceived visual forest characteristics identified by participants in the open visual sorts were vegetation density, light conditions, forest structural attributes and colours. The main perceived acoustic forest characteristics identified in the open acoustic sorts comprised bird song characteristics, physical properties such as volume, references to the time of day or seasonality and evoked emotions.

4. Perceived visual and acoustic diversity were significantly correlated with actual tree and bird species richness, respectively. Notably, the relationship was twice as strong for the acoustic sense. The acoustic sense may thus be crucial to obtain a more thorough understanding of perceived biodiversity.

5. We further computed several visual and acoustic diversity indices from the photos and audio recordings, e.g. colourfulness or acoustic complexity, and assessed their relevance for perceived and actual diversity. While all acoustic diversity indices were significantly associated with perceived acoustic diversity and bird richness, we could not identify a visual diversity index that captured perceived visual diversity and tree richness.

6. Our results suggest that people can perceive species richness. Our identified visual and acoustic forest characteristics may help to better understand perceived diversity and how it differs from actual diversity. We present acoustic diversity indices that quantify aspects of perceived and actual acoustic diversity. These indices may serve as cost-efficient tools to manage and plan greenspaces to promote biodiversity and mental well-being.

## 1. Introduction

Amidst a global biodiversity and health crisis, a growing body of research identifies the importance of biodiversity for people, on both a global level, for example with regards to ecosystem service provisioning, and the individual level, relating to one’s health and well-being (e.g. Marselle et al., 2019; Marselle et al., 2021). We are, however, not only experiencing an extinction of species but also an extinction of biodiversity experience due to rural-urban migration and less nature contact (Soga & Gaston, 2016). A better understanding of aspects of biodiversity that people predominantly perceive may provide leverage points to foster people’s interaction with biodiversity. This understanding could then help inform natural management strategies that benefit humans and biodiversity conservation (e.g. Pritchard, 2021; Reinecke & Blum, 2018).

There is a consensus across studies that green spaces can provide health-promoting effects (Bowler et al., 2010). For example, visits to the forest directly increase mental health and well-being (Rozario et al., 2024), while indirectly fostering physical health since forests can buffer heat stress (Gillerot et al., 2022; 2024) or improve air quality (Smith et al., 2013; Steinparzer et al., 2022). However, the potential incremental value of biodiversity of such green spaces to health is less understood. A growing research area is working to more finely establish the links between biodiversity and mental health and well-being (hereafter mental well-being; Marselle et al., 2019; Hedin et al., 2022; Lovell et al., 2014). In a large-scale, pan-European study Methorst et al. (2021a) found that mental well-being was significantly positively associated with bird species richness, with the effect of an increase in bird species richness comparable to an increase in income. Bird species richness was also positively associated with greater mental well-being in an epidemiological study of Germany (Methorst et al., 2021b). For forest ecosystems, Wolf et al. (2017) showed that watching videos of forests with four opposed to one tree species positively influenced mental well-being, while Rozario et al. (2024) and Nghiem et al. (2021) found that greater perceived levels of biodiversity in forests were associated with increased mental well-being.

Findings for biodiversity and mental well-being, however, are mixed, due to variations in study designs and how biodiversity and mental well-being are measured (e.g. Marselle et al., 2019). Biodiversity has different facets as it encompasses variations on the genetic, organismic and ecological level (Heywood & Watson, 1995), which can be described on multiple spatial and temporal scales, and can be further categorised into compositional, functional and structural diversity (Noss, 1990). Thus, there is no single measure of biodiversity and each measure is only a proxy for the “true” biodiversity it is supposed to describe. In forest ecosystems, biologists and forest managers alike therefore rely on a number of measurements, – and indices derived from those measurements – to aid them in quantifying biodiversity. To describe forest diversity, structural and organismic diversity is often considered. Structural diversity can be quantified by a variety of measures such as diameter at breast height, tree height, basal area (density), canopy layers or leaf area (e.g. Maes et al., 2011; Storch et al., 2018; Van Loy et al., 2003). Organismic diversity, on the other hand, can be quantified by tree and understorey plant species richness, but also by the diversity of animal species that rely on the forest habitat for one or more stages of their life cycles, such as forest bird or invertebrate richness. Indices such as Shannon diversity are then used to assess how diverse a community a given habitat is able to support. Indices can also measure attributes of a habitat aside from organismic diversity, such as remotely sensed structural indices derived from Light Detection and Ranging (LiDAR) measurements (e.g. the Stand Structural Complexity Index (SSCI, Ehbrecht et al., 2017; Frey et al., 2019). The complexity of sound emanating from a forest habitat can also be measured using e.g. the Acoustic Complexity Index (ACI; Pieretti et al., 2011). Ecological indices provide a measurement of one or more characteristics of a feature being studied, be that organismic diversity, stand structural complexity or acoustic complexity to help quantify an aspect of biodiversity.

To date, studies investigating the effect of forest diversity on mental well-being have been limited in the number of measures used to represent biodiversity, and often select a single (and differing) measure for each study (Grilli & Sacchelli, 2020; Hedin et al., 2022; Marselle et al., 2019). With even a single measure such as species richness, mental well-being responses can further vary depending on species traits and the mental well-being dimension assessed (An et al., 2019; Elsadek et al., 2019; Fisher et al., 2023; Guan et al., 2017; Sivarajah et al., 2018).

In fact, studies also differ in how biodiversity is conceptualised: measured or perceived (Marselle et al., 2021). A proxy measure of measured biodiversity, perceived biodiversity refers to subjective estimations of the biodiversity present in an environment (Dallimer et al., 2012; Fuller et al., 2007; Rozario et al., 2024). Lately, there has been a growing interest in establishing the link between measured and perceived biodiversity to determine the accuracy of this proxy measure. Most studies confirm a relationship between both (Ferraro et al., 2020; Fuller et al., 2007; Gao et al., 2019; Gonçalves et al., 2021; Johansson et al., 2014; Lindemann-Matthies et al., 2010; Rozario et al., 2024; Southon et al., 2018) but associations are often weak (Rozario et al., 2024) or non-existent (Dallimer et al., 2012; Phillips & Lindquist, 2021; Stobbe et al., 2022). This indicates that, despite a substantial overlap, there is a divergence between biodiversity measured by biologists and how people perceive it.

Studies have also juxtaposed the associations of measured and perceived biodiversity with mental well-being. In these cases, perceived biodiversity was found to have a stronger effect on mental well-being than measured biodiversity (Cameron et al., 2020; Dallimer et al., 2012; Farris et al., 2024; Rozario et al., 2024; Schebella et al., 2019; Zumhof, 2019). Indeed, Rozario et al. (2024), Farris et al. (2024) and Zumhof (2019) only found significant mental well-being effects for perceived biodiversity. A more nuanced differentiation between measured and perceived biodiversity is hence needed to understand their respective effects on mental well-being.

Importantly, perceiving biodiversity is a multisensory experience (Franco et al., 2017; Hedblom et al., 2019). Given the disproportionate amount of studies focused on the visual sense, a better understanding of the contribution of other senses (individually and combined) to perceiving biodiversity – and its effects on mental well-being – is desirable. Fisher et al. (2023), for instance, found that sounds were among the most frequently recognised forest diversity characteristics, alongside visual cues such as colour, and that sounds elicited the greatest well-being responses. Higher levels of acoustic diversity, in terms of vocalising bird species, have further been shown to reduce symptoms of depression (Stobbe et al., 2022) and elicit more pronounced restorative effects (Ferraro et al., 2020; Uebel et al., 2021). The acoustic sense in particular may thus be crucial to obtain a more thorough understanding of perceived biodiversity.

For the visual sense, several studies investigated potential drivers of perceived diversity (Gonçalves et al., 2021; Hoyle, 2020; Hoyle et al., 2018; Southon et al., 2018). Hoyle et al. (2018) identified flower colour as a significant predictor of perceived plant richness, i.e. colourful meadows were perceived as more biodiverse. Southon et al. (2018) report higher perceived diversity ratings for meadows that were perceived as more colourful but also for meadows with greater vegetation height and evenness. Tree evenness and fruit showiness, however, were negatively associated with perceived diversity in urban parks, while positive links were found with species richness, butterfly evenness, leaf shape and evergreen species (Gonçalves et al., 2021). For the acoustic sense, no study, to date, has investigated potential drivers of perceived acoustic diversity.

Human perception involves both bottom-up processing of sensory information and top-down processing involving higher level mental functions, such as memory of past experiences, cognition (e.g. knowledge, expectations, attention, motivation) and emotions (Axelrod, 1973; Brunswik, 1952; Friston, 2005; Gregory, 1980; Koffka, 1922). It is these top-down processes that determine what is ultimately perceived (e.g Axelrod, 1973). Perceived biodiversity therefore is the consequence of how biophysical properties such as colour for the visual sense or sound pressure level for the acoustic sense are translated into sensory input and how this sensory input is further processed and interpreted by the mental system.

With a focus on the visual and acoustic senses, we aim to identify biophysical and subjective factors that influence perceived biodiversity. We further investigate how perceived visual and acoustic diversity relate to species richness and established ecological diversity indices. Our specific research questions are:

1. What do people perceive when seeing or hearing different levels of biodiversity and what mental representations do they employ?
2. How congruent are perceived biodiversity and measured species richness (‘actual diversity’)?
3. Are there diversity indices to quantify both perceived and actual diversity?

We test this using photographs and audio recordings of forest environments. In essence, limited to two senses, we assess what people experience in a forest, and if we can measure it. Based on this understanding, if we can identify diversity indices that simultaneously quantify a forest’s species richness and people’s perceptions of it, we can use these indices towards managing natural spaces both to increase biodiversity and their experienced value that could foster mental well-being.

## 2. Methods

We conducted sorting experiments (Chollet et al., 2014; Lobinger & Brantner, 2020) to make mental representations associated with forest diversity tangible (Austen et al., 2021, 2023). Participants sorted forest photos (‘visual sort’) and audio recordings (‘acoustic sort’) of varying species richness, to identify subjective visual and acoustic forest characteristics that stood out to them and to obtain perceived visual and acoustic diversity ratings (Fig.1, Step 1) that we then correlated with species richness (Fig.1, Step 2). Next, we computed diversity indices from the photographs and recordings and tested if these indices were correlated with actual diversity (tree and bird richness) and perceived diversity (Fig.1, Step 3).

**Figure 1.**
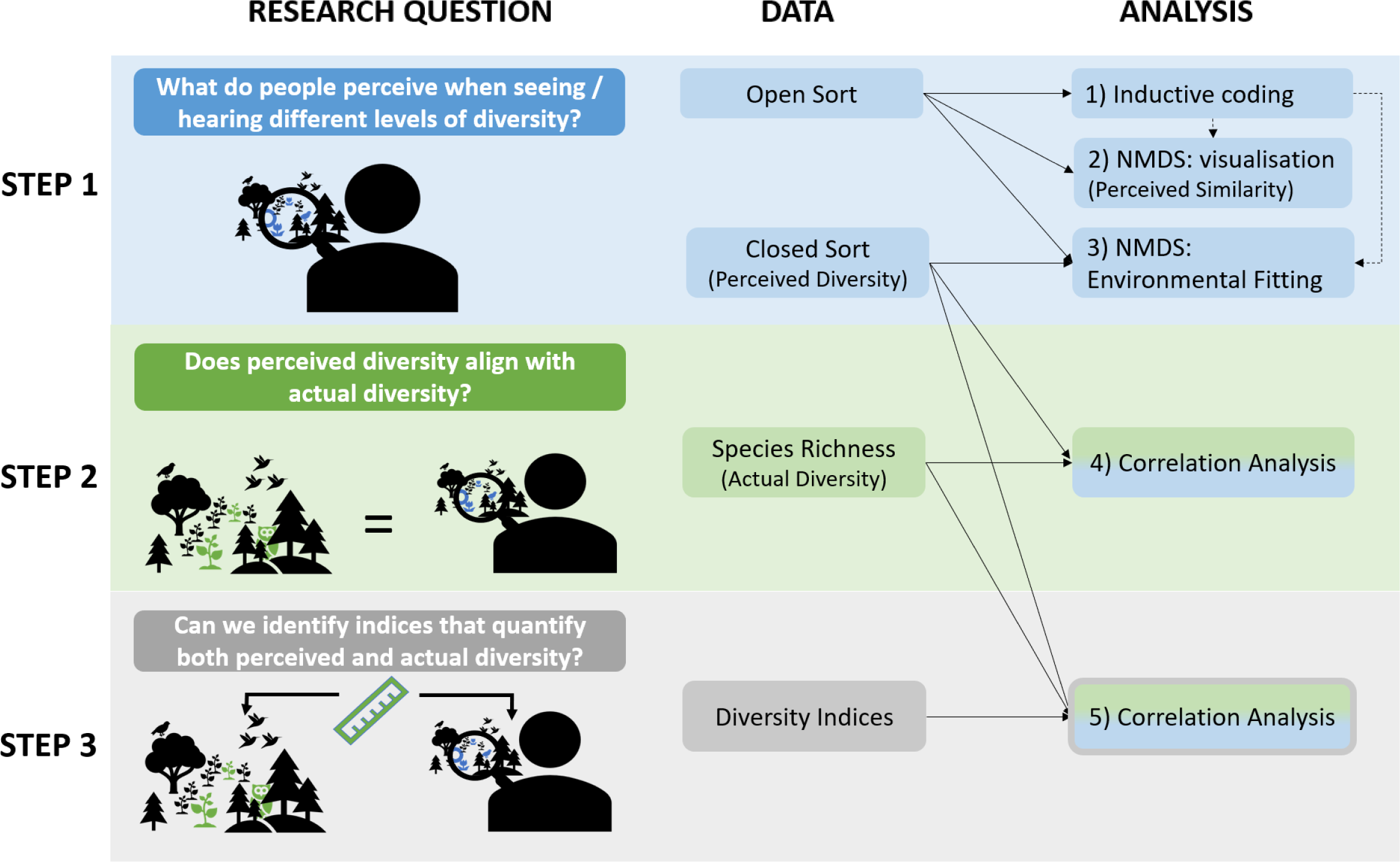
Conceptual figure of the study objectives. NMDS = non-metric multidimensional scaling. Solid arrows indicate that data was used for respective analyses. Dashed arrows show the conceptual relationship between the sorting criteria and the NMDS plots / environmental fitting.

### 2.1 Stimulus Materials

The stimulus set for the visual sort comprised 57 pre-selected forest photographs depicting varying levels of tree species richness (1-5 species), including varying combinations of tree species within each richness category (see Appendix A1 for detailed information about the photographs). Photographs were taken in temperate forests managed with close-to-nature forestry schemes in Germany (Hainich), Belgium (TREEWEB forest patches, https://treedivbelgium.ugent.be/pl_treeweb.html) and Poland (Białowieża) in late summer 2020. Forest patches were pre-identified and categorised based on tree richness by forest ecologists and local forestry agencies. Photos were taken under consistent weather and time conditions (cloudy to sunny / 11am-4pm; see Bergen et al., 1995; Carrus et al., 2015; Hofmann et al., 2017), with an iPhone 6s (12-megapixel camera) and a polarisation filter in order to parcel out scattered light that reduces image quality. We further took the photos at a height of ∼ 1.80 metres to ensure a realistic human observer perspective (Grassini et al., 2019). Photos were printed out in A5 format and laminated for multiple use.

The stimulus set of the acoustic sort consisted of 16 ten-second long (Ratcliffe et al., 2016, 2020) natural audio recordings taken from temperate forests (see Appendix A2 for detailed information about the audio recordings). The primary gradient represented in these recordings was vocalisations from varying numbers of unique bird species (bird species richness), from zero to six bird species in a given recording. We used fewer stimuli than in the visual sort as in acoustic sorts, recordings have to be listened to sequentially. It would have thus been overly cognitively demanding and time-consuming for participants to remember (and re-play) 57 sound clips in order to complete the task (see e.g. Berland et al., 2015; Giordano et al., 2011; Maffiolo et al., 1998 for numbers of audio recordings in other acoustic sorts).

### 2.2. Participants

For the visual sort, we tested 48 participants (41 women, 85%) between 18 and 35 years of age (*M* = 23.92, *SD* = 4.20). While reasonable sorting clusters can be obtained with 20-30 participants (Chollet et al., 2014; Harloff & Coxon, 2007; Rugg & McGeorge, 2005; Tullis et al., 2004), we chose to work with a higher number of participants, as the robustness of results in sorting studies increases with the number of participants (Berland et al., 2015; Blancher et al., 2012). Robustness further increases as a function of participants’ expertise with the stimuli to be sorted and decreases with higher task complexity (Blancher et al., 2012). A majority of the study participants were psychology students, thus with little expertise in ecology or forestry. In addition, we assumed that the complexity of the visual sort was considerably high as (i) the content of the photographs was similar which impedes the clustering of photos in the sorting process and (ii) the number of photos exceeded recommendations (see Chollet et al., 2014; Rugg & McGeorge, 2005) which poses an increased demand on working memory capacity. For the acoustic sort, we also tested 48 participants (34 women, 71%) between 18 and 35 years of age (*M* = 24.33, *SD* = 4.45) to achieve an equivalent sample size.

Prerequisites for study participation were good general health as well as normal or corrected-to-normal vision (visual sort) and hearing (acoustic sort). Participants for both studies were recruited through the mailing list of the Cognitive and Biological Psychology group of the Wilhelm Wundt Institute for Psychology at Leipzig University, various social media platforms and word of mouth. The procedure for both studies followed the principles of the Declaration of Helsinki. Ethical approvals were obtained from the local ethics committee at Leipzig University (reference number visual sort: 2021.02.26_eb_77; reference number acoustic sort: 2022.07.07_eb_163). All participants gave written informed consent prior to participation. There was no financial remuneration; psychology students (visual sort: *n*=24; acoustic sort: *n*=27) received course credits that equalled the time spent at the experiment in hours.

### 2.3 Procedure

Data collection took place from March to May 2021 (visual sort) and August to November 2022 (acoustic sort) in facilities of Leipzig University and the German Centre of Integrative Biodiversity Research (iDiv).

For both the visual and acoustic sort, we combined two, single-criterion open sorts with one closed sort (see Canter, 1996; Chollet et al., 2014; Harloff & Coxon, 2007; Rugg & McGeorge, 2005 for different sorting paradigms). In open sorts, participants were free to sort objects according to any similarity-based sorting criterion they like. For example, a participant could choose to sort the forest photos based on sorting criteria, such as vegetation density or colour. They then choose subordinate categories to assign the photos to, e.g. along a gradient such as low, medium and high vegetation density. In closed sorts, participants are asked to sort objects based on pre-established criteria. In the closed visual sort, we asked participants to sort the photographs according to low, medium and high forest diversity (Phillips & Lindquist, 2021; White et al., 2017). In the closed acoustic sort, we asked participants to sort the audio recordings according to low, medium and high acoustic forest diversity.

For the visual sorts, participants were instructed to distribute the 57 photos on either the floor or tables in a way that enabled parallel viewing of all photos. Sound recordings for the acoustic sorts were digitally presented on the desktop of a Dell Latitude E7440. Participants listened to the recordings with headphones (HyperX Cloud Stinger™ Core) and loudness was set to volume level 34 locally on the laptop. We instructed the participants to listen to all 16 sound recordings at least once but as often as they liked before and while sorting.

In both experiments, participants began with the two open sorts as conducting the closed sort prior to the open sorts would have potentially primed the sorting behaviour in the open sort towards diversity. We conducted two open sorts to obtain more visual and acoustic forest characteristics from the participants and to increase the robustness of the results (Blancher et al., 2012). The instructions for the open sorts were identical for the open visual and acoustic sorts and for the two open sorts conducted per sense, with the restriction that two distinct sorting criteria had to be employed (e.g. “vegetation density” for the first open visual sort and “colour” for the second open visual sort). For each open sort, participants had to identify at least two subordinate categories (e.g. low and high vegetation density). At least one sorting object had to be assigned to each sub-category.

Sorting criteria and sub-categories, as well as the assignment of photos and audio recordings to sub-categories were recorded on documentation sheets. Numbers on the back of each photograph as well as labels for the audio files on the desktop were used to track the assignment of stimuli to sub-categories. Results were captured in paper-pencil format. The time of data collection varied between 1.5-3.5 h for the visual sorts and between 1 and 1.5 h for the acoustic sorts since no time restriction was given.

### 2.4 Diversity Indices

We wanted to test if we could compute diversity indices from the stimuli (photos/recordings) that measured actual diversity (i.e. species richness) and perceived diversity (see Fig. 1, Step 3). Four visual indices were selected to reflect diversity related to characteristics of photographs that participants might observe, such as the intensity of green, colour as proxy for vegetation biomass and variations in light (Frey et al., 2019; Hasler & Süsstrunk, 2003; Menzel & Reese, 2021). We also computed four acoustic indices known in prior studies as reliable indicators of biodiversity (Alcocer et al., 2022; Yip et al., 2022). These indices quantify different characteristics of an acoustic recording, for example the complexity of a bird vocalisation, density of calls within a recording, or the frequency bands that birds vocalise in (e.g. low-frequency pigeon ‘coos’ versus higher-frequency warbles). Indices were selected to also match an accompanying study where respective indices were related to forest features (Gillerot et al., in preparation). See Table 1 for a list of the computed indices, Appendix A3 for summary statistics and code used to compute them and Appendix A4 for correlations between indices.

**Table 1.**
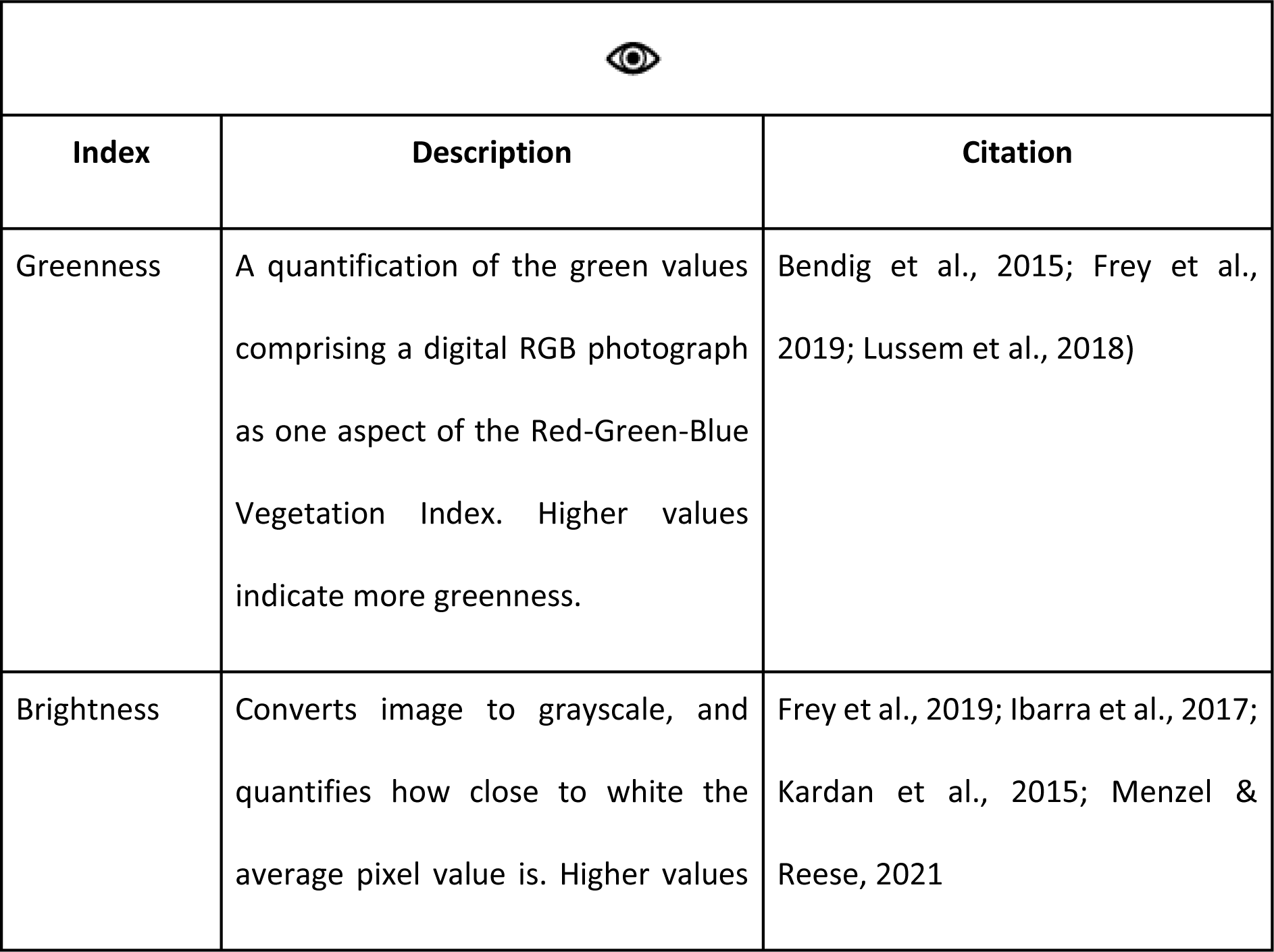
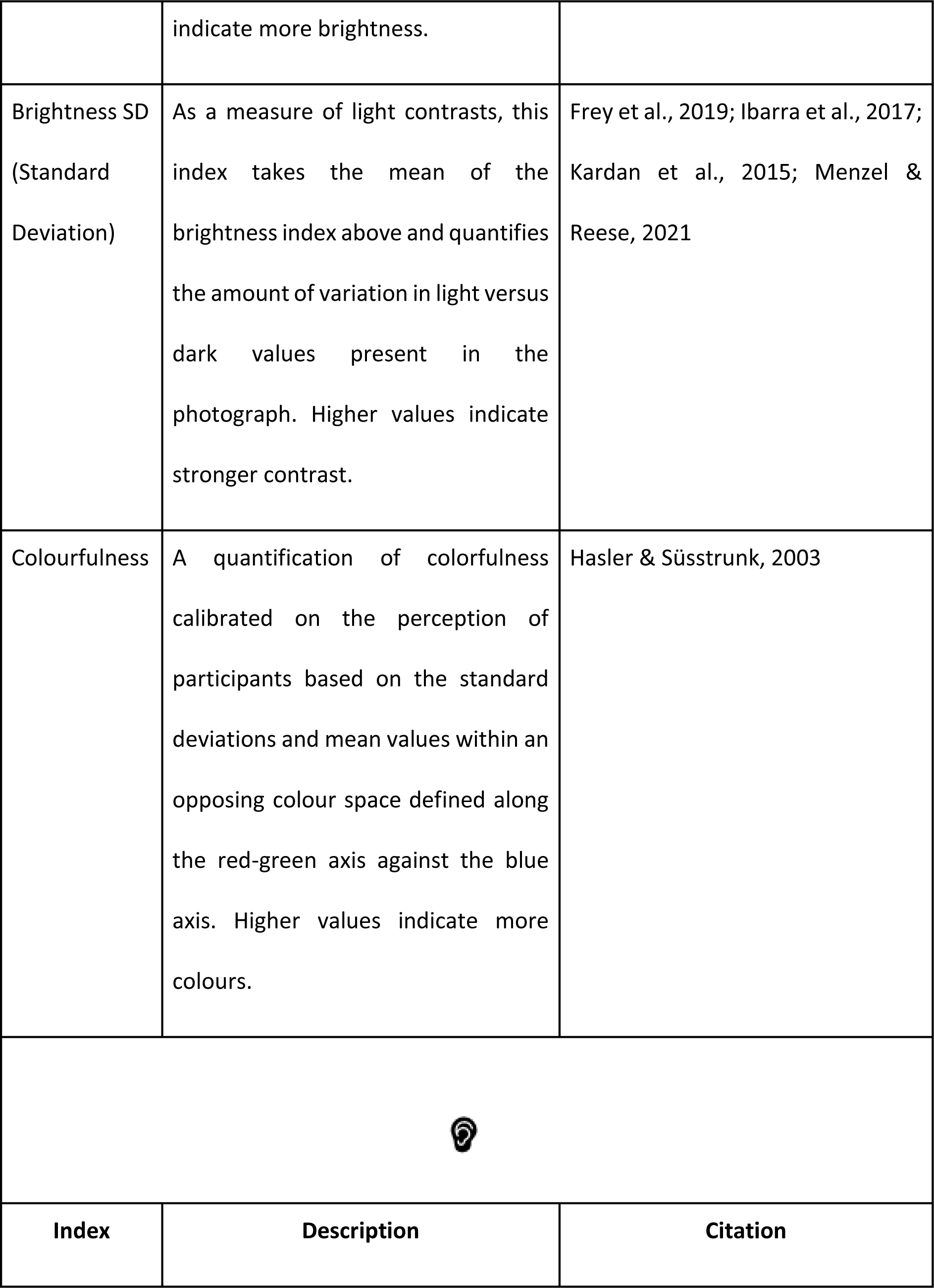
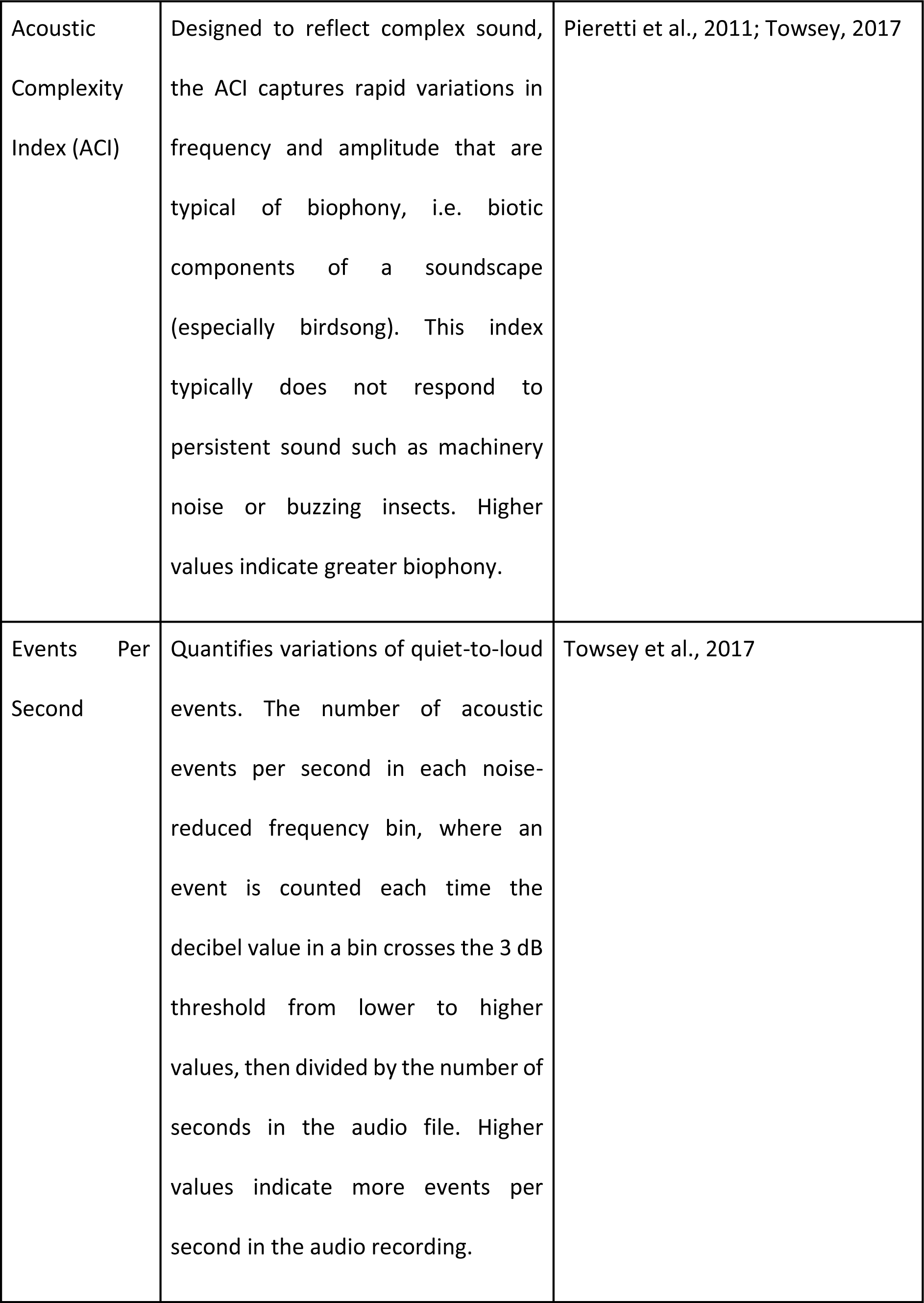
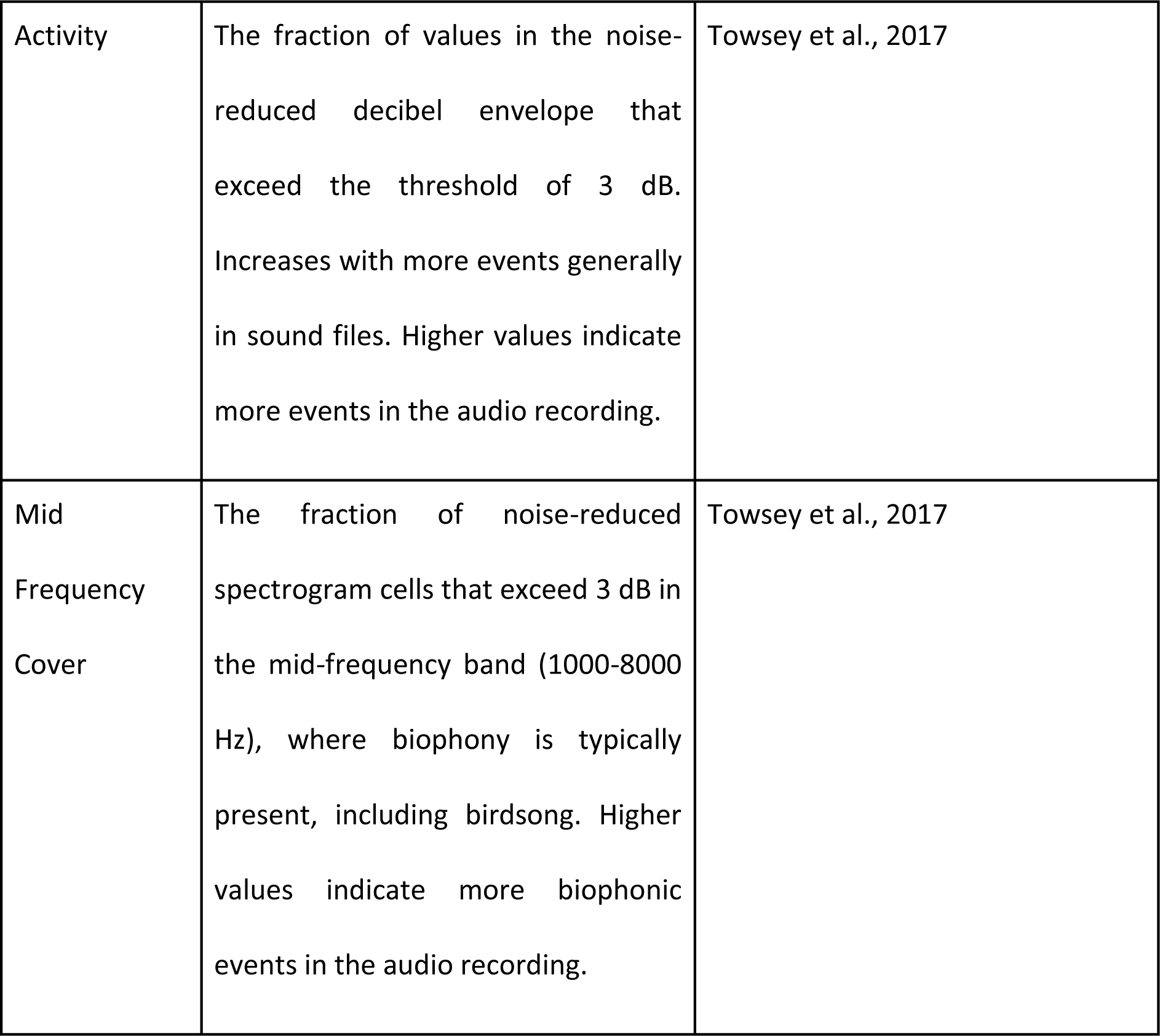
Computed indices on visual and acoustic stimuli.

|  |  |  |
| --- | --- | --- |
| Index | Description | Citation |
| Greenness | A quantification of the green values comprising a digital RGB photograph as one aspect of the Red-Green-Blue Vegetation Index. Higher values indicate more greenness. | Bendig et al., 2015; Frey et al., 2019; Lussem et al., 2018) |
| Brightness | Converts image to grayscale, and quantifies how close to white the average pixel value is. Higher values | Frey et al., 2019; Ibarra et al., 2017; Kardan et al., 2015; Menzel & Reese, 2021 |
|  | indicate more brightness. |  |
| Brightness SD<br>(Standard Deviation) | As a measure of light contrasts, this index takes the mean of the brightness index above and quantifies the amount of variation in light versus dark values present in the photograph. Higher values indicate stronger contrast. | Frey et al., 2019; Ibarra et al., 2017; Kardan et al., 2015; Menzel & Reese, 2021 |
| Colourfulness | A quantification of colourfulness calibrated on the perception of participants based on the standard deviations and mean values within an opposing colour space defined along the red-green axis against the blue axis. Higher values indicate more colours. | Hasler & Süssstrunk, 2003 |
| **Index** | **Description** | **Citation** |
| Acoustic Complexity Index (ACI) | Designed to reflect complex sound, the ACI captures rapid variations in frequency and amplitude that are typical of biophony, i.e. biotic components of a soundscape (especially birdsong). This index typically does not respond to persistent sound such as machinery noise or buzzing insects. Higher values indicate greater biophony. | Pieretti et al., 2011; Towsey, 2017 |
| Events Per Second | Quantifies variations of quiet-to-loud events. The number of acoustic events per second in each noise-reduced frequency bin, where an event is counted each time the decibel value in a bin crosses the 3 dB threshold from lower to higher values, then divided by the number of seconds in the audio file. Higher values indicate more events per second in the audio recording. | Towsey et al., 2017 |
| Activity | The fraction of values in the noise-reduced decibel envelope that exceed the threshold of 3 dB. Increases with more events generally in sound files. Higher values indicate more events in the audio recording. | Towsey et al., 2017 |
| Mid Frequency Cover | The fraction of noise-reduced spectrogram cells that exceed 3 dB in the mid-frequency band (1000-8000 Hz), where biophony is typically present, including birdsong. Higher values indicate more biophonic events in the audio recording. | Towsey et al., 2017 |

### 2.5 Statistical Analyses

#### 2.5.1 Perceived visual and acoustic forest characteristics

We qualitatively analysed the sorting criteria reported by participants in the open sorts, to better understand the perceived forest characteristics that potentially influenced perceived diversity (Fig.1 Step 1, Analysis 1). In total, 92 open visual sorts and 96 open acoustic sorts were completed and analysed separately. Overarching clusters within the sorting criteria were identified following a general inductive approach, i.e. clusters were formed in a bottom-up manner based on the precise wording of the sorting criteria given by participants (Thomas, 2006). To increase objectivity, we conducted independent parallel coding (Thomas, 2006) with five raters from various disciplines (KR (psychology), MM (psychology), RRYO (ecology), MGB (philosophy), and OS in acknowledgements (neurosciences)), the results of which we refer to as first-order clusters. Raters could form as many first-order clusters as they considered appropriate to cover the variety of themes they identified in the sorting criteria. At least one first-order cluster had to be assigned to each sorting criterion. First-order clusters were then further summarised by one rater (KR) to form the final set of eight perceptual visual and acoustic forest characteristics (Thomas, 2006), hereafter referred to as second-order clusters.

#### 2.5.2 Perceived visual and acoustic forest similarity

All of the following analyses were computed in R Statistical Computing Environment (version 4.2.1; R Core Team, 2022). We applied non-metric multidimensional scaling (NMDS) on the open sort data to assess how similar the photographs and recordings were perceived in relation to one another (Fig.1, Step 1, Analysis 2). Similarity or co-occurrence matrices of the sizes 57×57 (open visual sort) and 16×16 (open acoustic sort) were calculated by comparing pairs of photos or recordings for each participant. For each pair, we counted the number of common assignments to sub-categories. These counts were then used to populate the corresponding cells in the matrices (see Fig. 2 for a schematic overview of the procedure). We then calculated Canberra distances (Gerstenberg & Hofmann, 2016) to convert the similarity matrices into dissimilarity matrices as Canberra distances are well-suited for non-standardised count data and particularly robust to outliers (Roberts, 2017). Afterwards, we computed an NMDS analysis using the *vegan* package (Oksanen et al., 2007). We selected the number of appropriate NMDS dimensions based on stress values (below 0.1; Kruskal, 1964). Two NMDS dimensions produced stress values of 0.06 for the visual sort and 0.04 for the acoustic sort indicating fair-to-good fit of the data to reduced ordination space.

**Figure 2.**
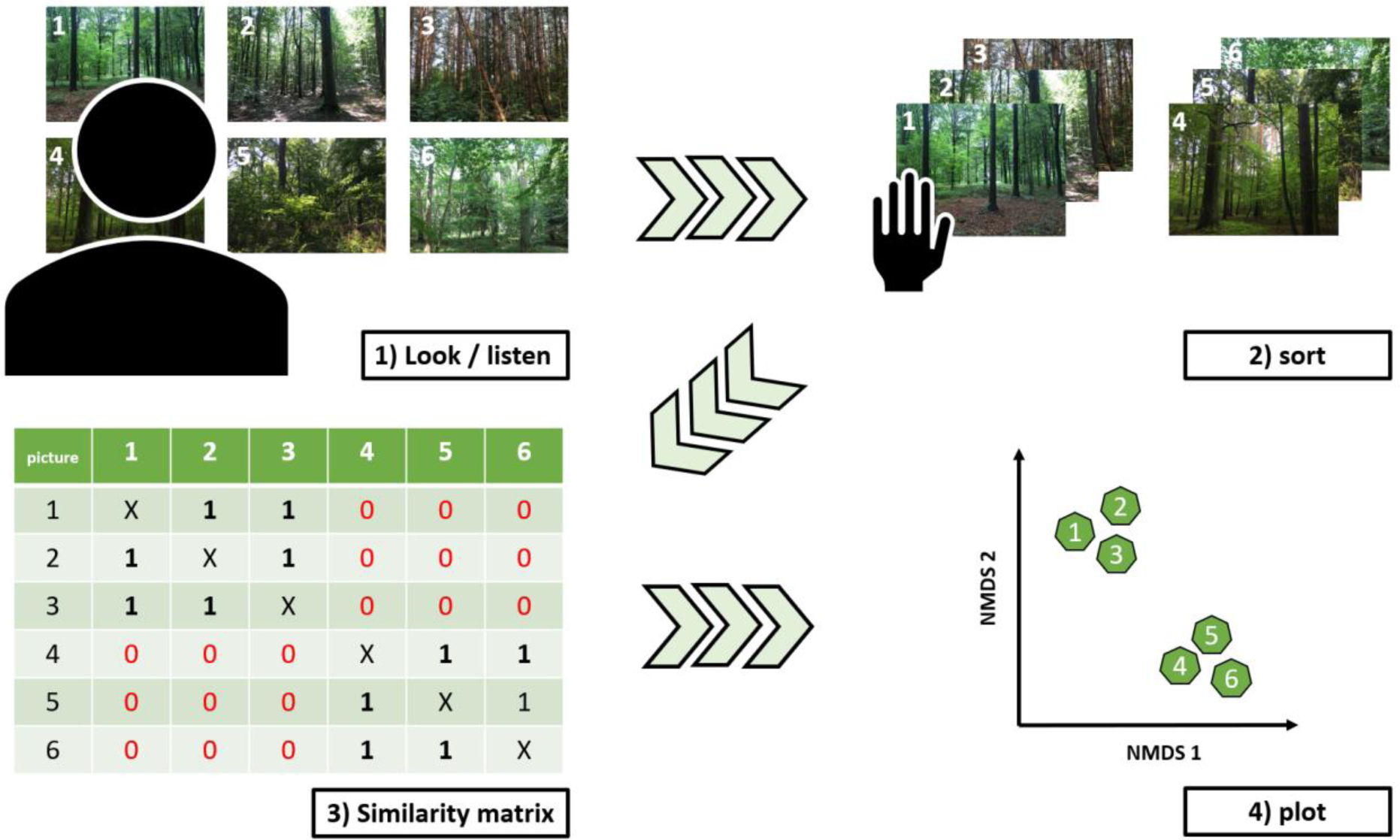
Schematic overview of the non-metric multidimensional scaling (NMDS) procedure using the open visual sort as an example. Each cell of the similarity matrices contained count values of common assignments of pairs of photos/recordings to categories. In this example, photos 1,2 and 3 have been assigned to the same category, so a ‘1’, representing one count, appears at the intersection of respective photos in the similarity matrix.

#### 2.5.3 Perceived visual and acoustic forest diversity

Perceived diversity ratings for the visual and acoustic stimuli were calculated using data from the closed sorts. The three levels of the closed sorts were coded as low diversity = 1, medium diversity = 2 and high diversity = 3. Mean perceived diversity scores were calculated for each photo and recording, respectively. To validate whether we could describe perceived diversity well with the open sort criteria, we conducted an environmental fitting analysis (Fig. 1, Step 1, Analysis 3). This enabled us to investigate whether perceived diversity explained the participants’ open sorting patterns, i.e. the produced NMDS solutions, using the ‘envfit’ function in the *vegan* package (Oksanen et al., 2007).

#### 2.5.4 Associations between perceived and actual diversity and diversity indices

Finally, we tested the correlation between perceived and actual diversity, i.e. species richness (Fig. 1, Step 2), and between the diversity indices with perceived and actual diversity (Fig. 1, Step 3). Pearson’s correlations were used for parametric data, and Spearman’s correlations were used for non-parametric data, both using the *stats* package (R Core Team, 2022).

## 3. Results

### 3.1 Perceived diversity

#### 3.1.1 Perceived forest characteristics

The five raters assigned 501 first-order clusters to the original sorting criteria of the open visual sort (see Appendix A5 for the five raters’ first-order clusters). Based on those first-order clusters, eight second-order clusters were identified comprising vegetation density, light conditions, structure, colour, diversity, emotions, the ground layer and tree physical features, in descending order.

For the acoustic sort, a total of 507 first-order clusters were assigned to the sorting criteria (A5). The eight second-order clusters identified were: bird song characteristics, physical properties of the sound recordings, time, emotions, bird abundance, bird species, vitality of vocalisations and landscape features in descending order. Tables 2 and 3 show descriptions, examples and frequencies of the second-order clusters for the visual domain and acoustic domain, respectively.

**Table 2.**
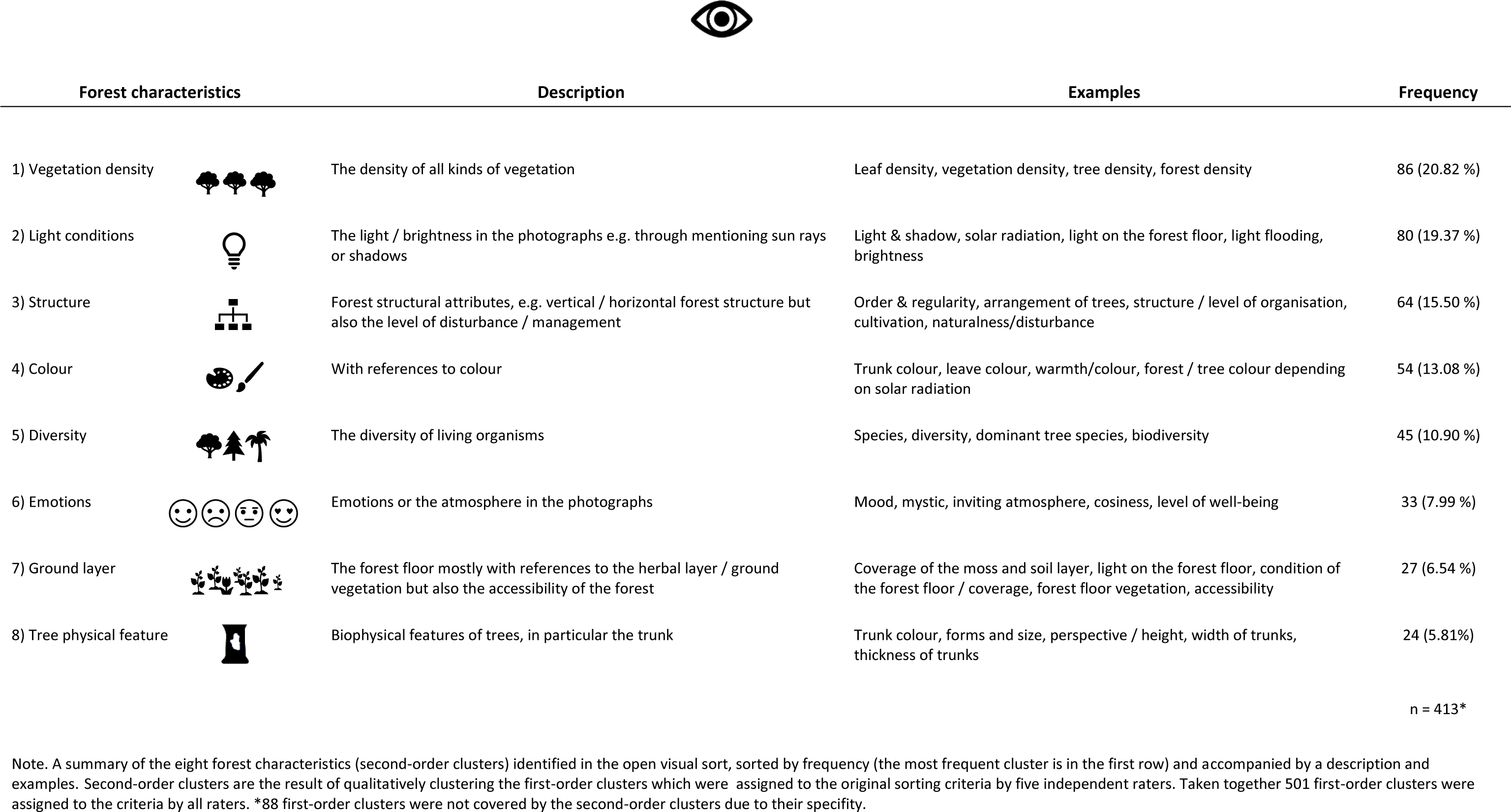
Visual forest characteristics identified in the open sorts.

| Forest characteristics |  | Description | Examples | Frequency |
| --- | --- | --- | --- | --- |
| 1) Vegetation density |  | The density of all kinds of vegetation | Leaf density, vegetation density, tree density, forest density | 86 (20.82 %) |
| 2) Light conditions |  | The light / brightness in the photographs e.g. through mentioning sun rays or shadows | Light & shadow, solar radiation, light on the forest floor, light flooding, brightness | 80 (19.37 %) |
| 3) Structure |  | Forest structural attributes, e.g. vertical / horizontal forest structure but also the level of disturbance / management | Order & regularity, arrangement of trees, structure / level of organisation, cultivation, naturalness/disturbance | 64 (15.50 %) |
| 4) Colour |  | With references to colour | Trunk colour, leave colour, warmth/colour, forest / tree colour depending on solar radiation | 54 (13.08 %) |
| 5) Diversity |  | The diversity of living organisms | Species, diversity, dominant tree species, biodiversity | 45 (10.90 %) |
| 6) Emotions |  | Emotions or the atmosphere in the photographs | Mood, mystic, inviting atmosphere, cosiness, level of well-being | 33 (7.99 %) |
| 7) Ground layer |  | The forest floor mostly with references to the herbal layer / ground vegetation but also the accessibility of the forest | Coverage of the moss and soil layer, light on the forest floor, condition of the forest floor / coverage, forest floor vegetation, accessibility | 27 (6.54 %) |
| 8) Tree physical feature |  | Biophysical features of trees, in particular the trunk | Trunk colour, forms and size, perspective / height, width of trunks, thickness of trunks | 24 (5.81%) |

n = 413\*
Note. A summary of the eight forest characteristics (second-order clusters) identified in the open visual sort, sorted by frequency (the most frequent cluster is in the first row) and accompanied by a description and examples. Second-order clusters are the result of qualitatively clustering the first-order clusters which were assigned to the original sorting criteria by five independent raters. Taken together 501 first-order clusters were assigned to the criteria by all raters. \*88 first-order clusters were not covered by the second-order clusters due to their specificity.

**Table 3.**
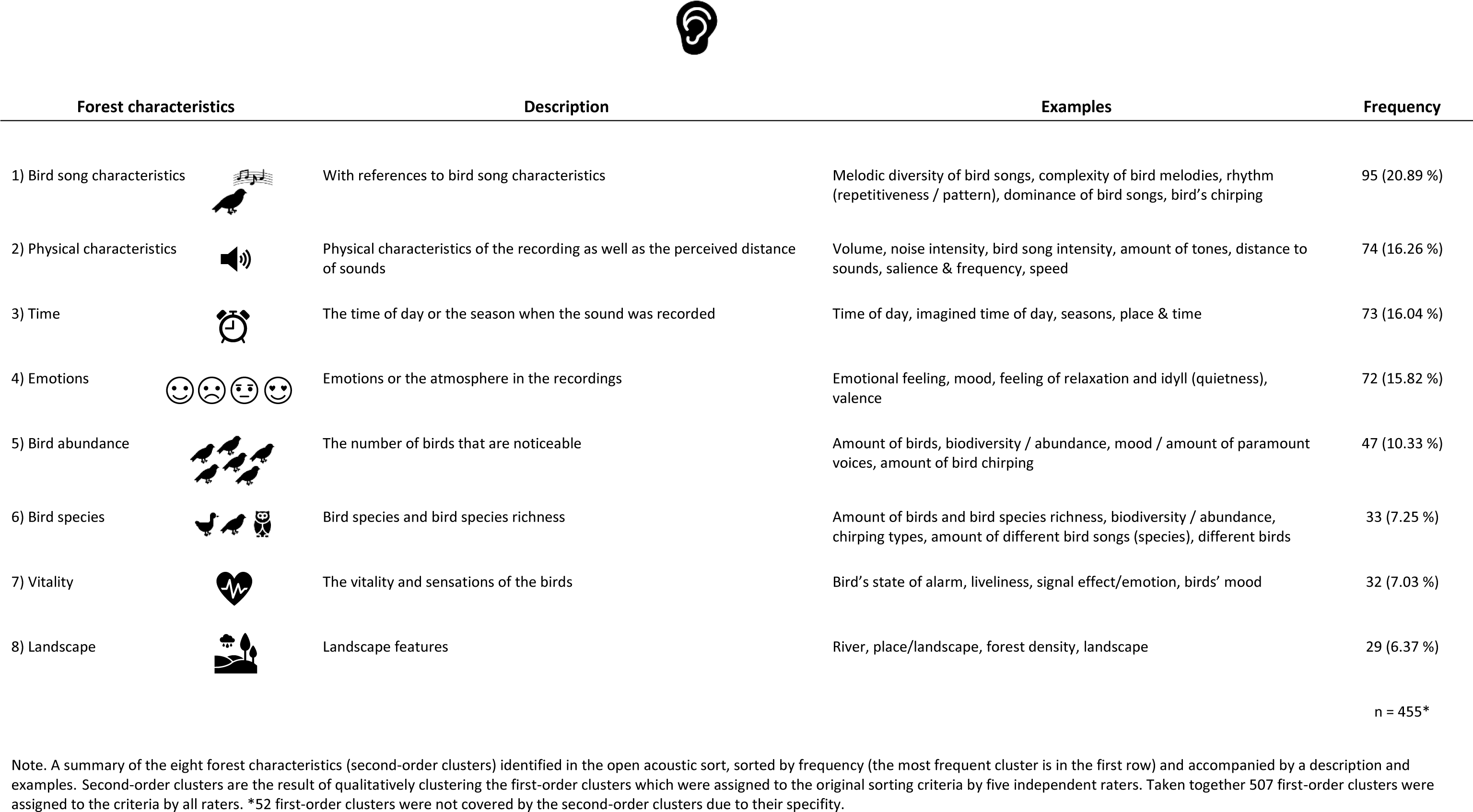
Acoustic forest characteristics identified in the open sorts.

| Forest characteristics |  | Description | Examples | Frequency |
| --- | --- | --- | --- | --- |
| 1) Bird song characteristics |  | With references to bird song characteristics | Melodic diversity of bird songs, complexity of bird melodies, rhythm (repetitiveness / pattern), dominance of bird songs, bird's chirping | 95 (20.89 %) |
| 2) Physical characteristics |  | Physical characteristics of the recording as well as the perceived distance of sounds | Volume, noise intensity, bird song intensity, amount of tones, distance to sounds, salience & frequency, speed | 74 (16.26 %) |
| 3) Time |  | The time of day or the season when the sound was recorded | Time of day, imagined time of day, seasons, place & time | 73 (16.04 %) |
| 4) Emotions |  | Emotions or the atmosphere in the recordings | Emotional feeling, mood, feeling of relaxation and idyll (quietness), valence | 72 (15.82 %) |
| 5) Bird abundance |  | The number of birds that are noticeable | Amount of birds, biodiversity / abundance, mood / amount of paramount voices, amount of bird chirping | 47 (10.33 %) |
| 6) Bird species |  | Bird species and bird species richness | Amount of birds and bird species richness, biodiversity / abundance, chirping types, amount of different bird songs (species), different birds | 33 (7.25 %) |
| 7) Vitality |  | The vitality and sensations of the birds | Bird's state of alarm, liveliness, signal effect/emotion, birds' mood | 32 (7.03 %) |
| 8) Landscape |  | Landscape features | River, place/landscape, forest density, landscape | 29 (6.37 %) |
|  |  |  |  | n = 455* |

Note. A summary of the eight forest characteristics (second-order clusters) identified in the open acoustic sort, sorted by frequency (the most frequent cluster is in the first row) and accompanied by a description and examples. Second-order clusters are the result of qualitatively clustering the first-order clusters which were assigned to the original sorting criteria by five independent raters. Taken together 507 first-order clusters were assigned to the criteria by all raters. \*52 first-order clusters were not covered by the second-order clusters due to their specificity.

#### 3.1.2 Perceived similarity

Environmental fitting indicated that perceived visual similarity, i.e. the sorting pattern in the open visual sorts, was significantly associated with perceived visual diversity in the closed sort (*p*<.001, *R^2^*=0.58; Fig.1, Step 1, Analysis 3). Perceived acoustic similarity was also significantly associated with perceived acoustic diversity (*p*<.001, *R^2^*=0.96), indicating that the open sorts (and the in 3.1.1 identified forest characteristics) represent aspects of perceived visual and acoustic diversity well (Fig.3, see A6 for the results of environmental fitting with perceived diversity, actual diversity and the diversity indices).

**Figure 3.**
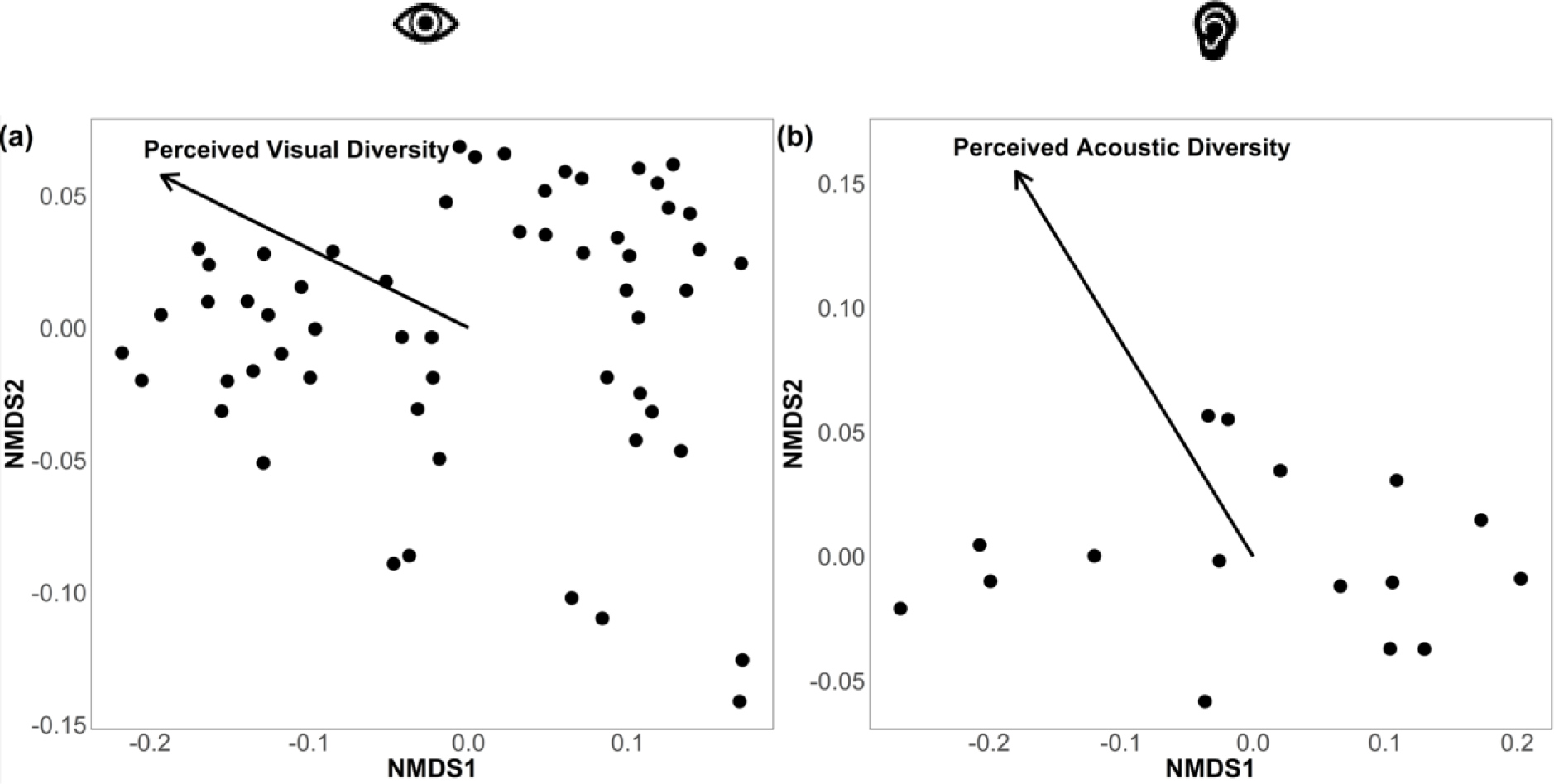
Non-metric multidimensional scaling (NMDS) plots resulting from the open visual (a) and open acoustic sort (b) (stress values: visual: 0.06; acoustic: 0.04). Points represent photos and recordings, respectively. The closer two points (shorter distance), the more similar photos or recordings were perceived as. Environmental fitting was conducted to see whether perceived diversity aligns with the produced NMDS solutions. The arrows illustrate the direction and strength of the associations between perceived diversity and the NMDS solutions.

### 3.2 Relationship between perceived and actual diversity

Positive significant correlations were found between perceived and actual diversity, i.e. species richness (Fig. 1, Step 2) for both the visual and acoustic stimuli (visual: Rho=.428, *p*<.001; acoustic: Rho=.869, *p*<.001; Fig. 4). A Fisher’s z-test was used to compare the strength of these two independent correlations using the ‘diffcor’ package (Blötner 2024). The correlation for the acoustic sense was significantly larger than the correlation for the visual sense (*z*=2.815; *p*=0.002; Cohen-q: 0.87), suggesting that participants were better in distinguishing different levels of bird richness compared to gradations of tree richness.

**Figure 4.**
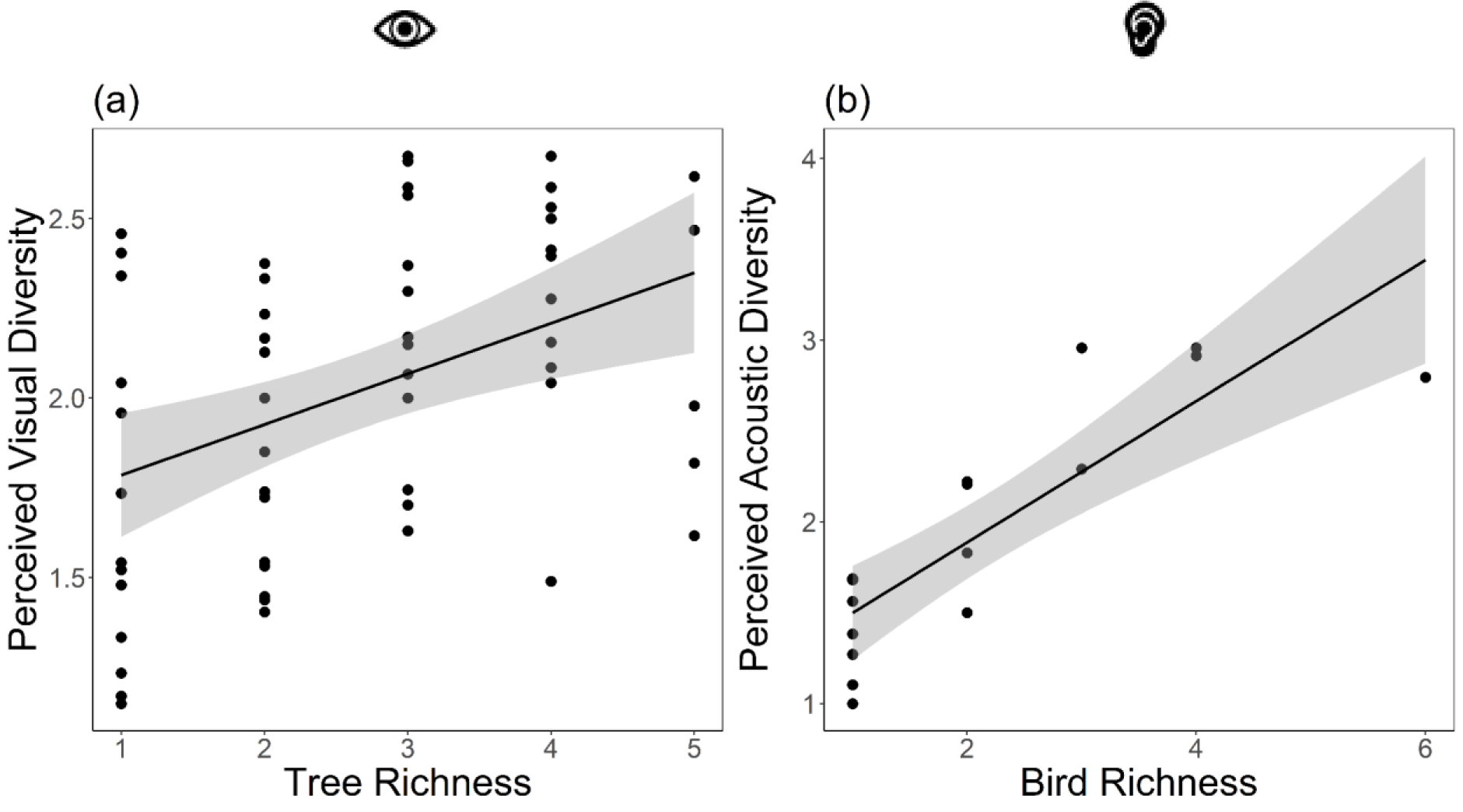
A linear regression to visualise the correlation between actual diversity (tree and bird species richness) and mean perceived visual (a) and acoustic (b) diversity per stimuli (57 photos and 16 audio recordings), as rated by participants. Shaded areas represent a 0.95 confidence interval.

### 3.3 Relationships between the computed diversity indices and perceived and actual diversity

Associations between the computed diversity indices and perceived diversity were mixed (Fig.1, Step 3; Table 4). Only one computed visual diversity index (Greenness) was significantly positively correlated with perceived visual diversity (Fig. 5a-d). All four computed acoustic indices, however, were positively significantly correlated with perceived acoustic diversity (Fig. 5e-h).

**Figure 5.**
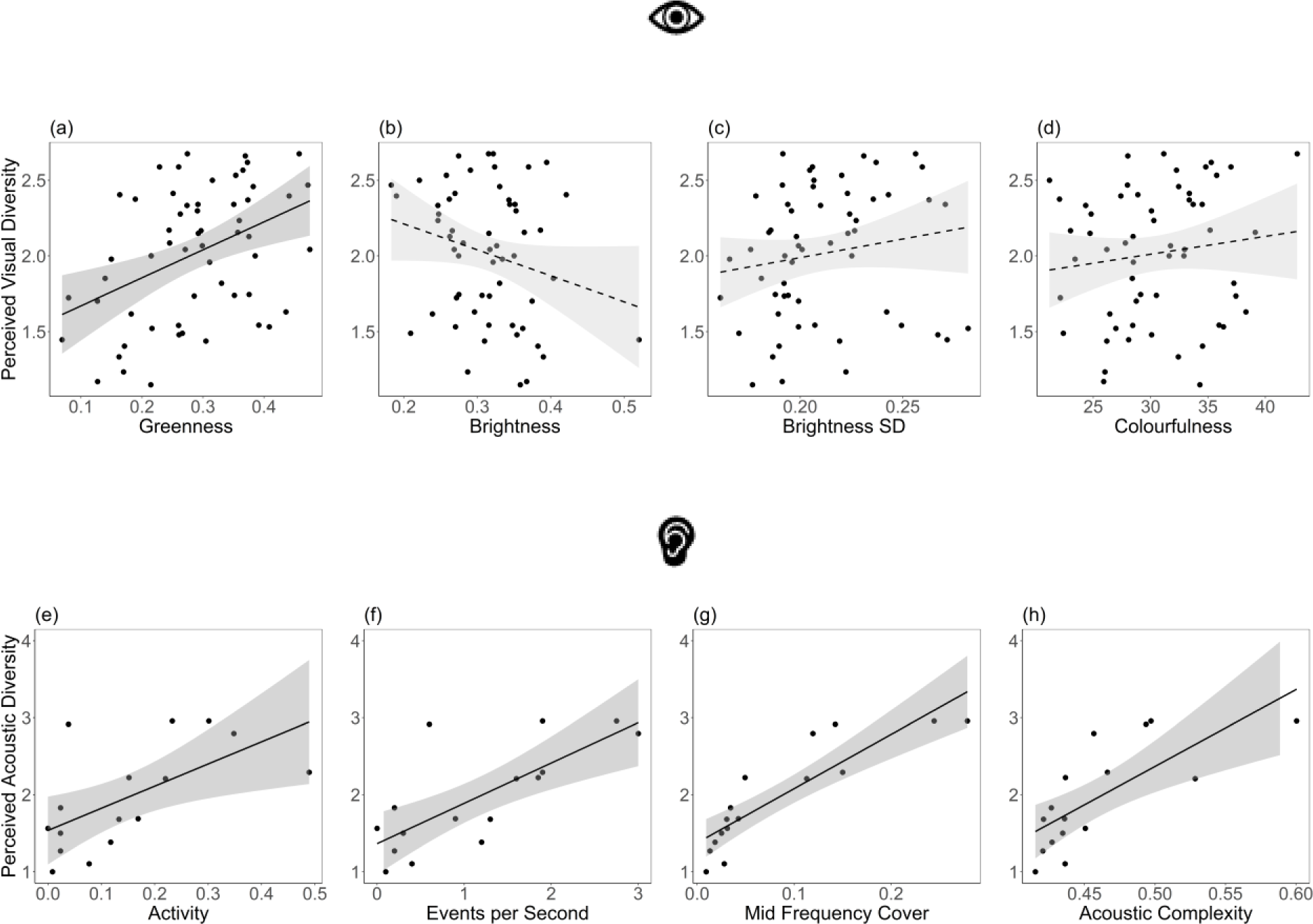
Correlations between the computed indices and perceived diversity. Panels a-d show correlations for visual indices derived from the photographs, and panels e-h represent acoustic indices derived from the audio recordings. Solid lines present significant correlations and dashed lines represent non-significant correlations, see details in Table 4. Shaded areas represent a 0.95 confidence interval.

**Table 4.**
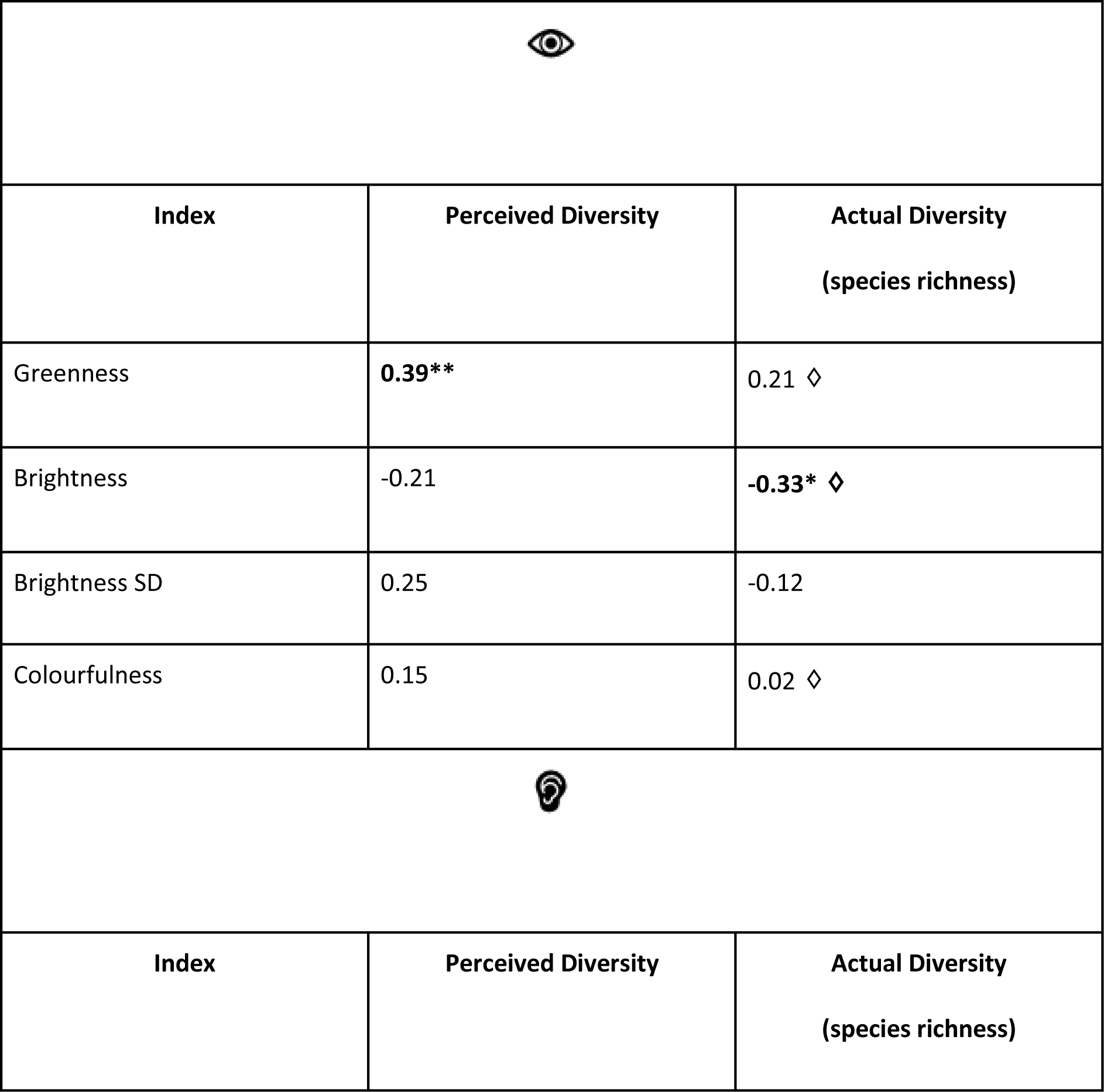
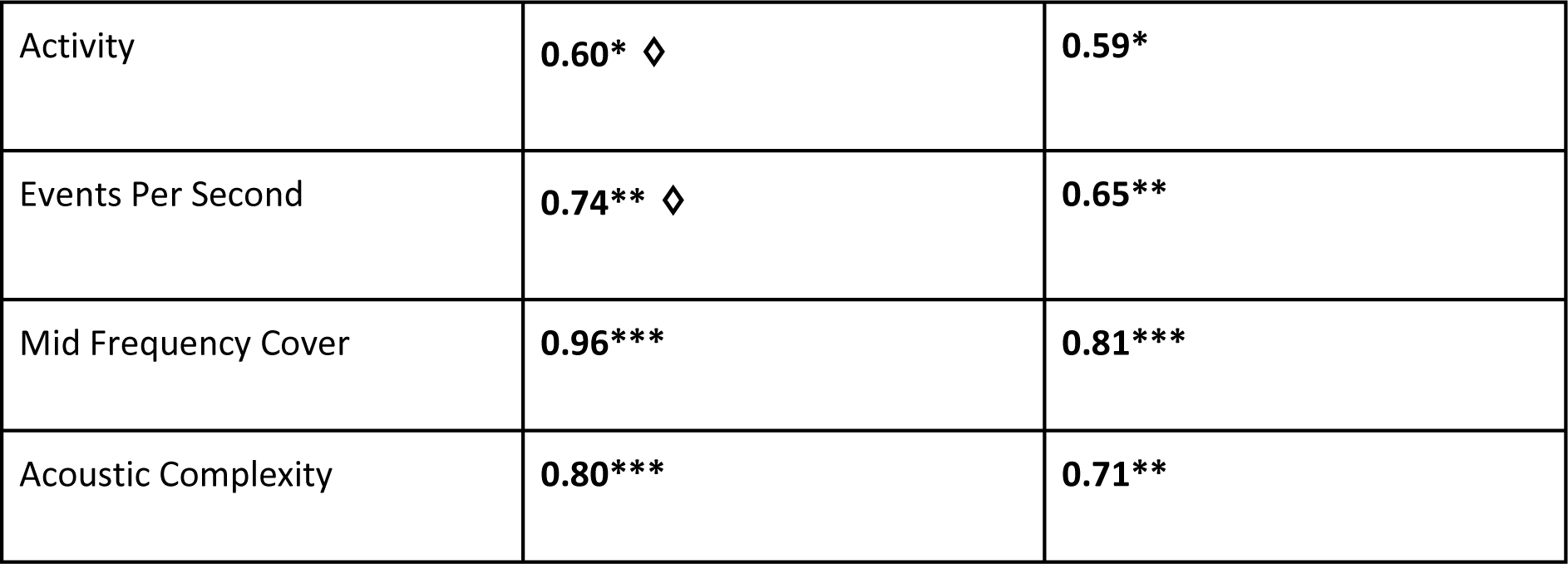
Correlation coefficients of comparisons between diversity indices, perceived diversity and actual diversity (tree and bird species richness). All tests were Spearman correlation tests, unless denoted by a ◊, indicating a Pearson correlation. Significant correlations are bolded and indicated as follows: < .05*, < .01**, <.001***.

|  |  |  |
| --- | --- | --- |
| Index | Perceived Diversity | Actual Diversity<br>(species richness) |
| Greenness | **0.39\*\*** | 0.21 $\diamond$ |
| Brightness | -0.21 | **-0.33\*** $\diamond$ |
| Brightness SD | 0.25 | -0.12 |
| Colourfulness | 0.15 | 0.02 $\diamond$ |
| Index | Perceived Diversity | Actual Diversity<br>(species richness) |
| Activity | **0.60\*** ◇ | **0.59\*** |
| Events Per Second | **0.74\*\*** ◇ | **0.65\*\*** |
| Mid Frequency Cover | **0.96\*\*\*** | **0.81\*\*\*** |
| Acoustic Complexity | **0.80\*\*\*** | **0.71\*\*** |

Finally, when comparing the computed diversity indices with actual diversity, i.e. species richness (Fig.1, Step 3), results followed a similar pattern to the comparison with perceived diversity (Table 4). Only the visual index brightness was significantly negatively correlated with tree species richness (Fig. 6a-d). This indicates that the stronger the grayscale values per pixels in a photo deviated from white, the less tree species the photo contained. All four acoustic indices were significantly positively correlated with actual acoustic diversity, i.e. bird species richness (Fig. 6e-h).

**Figure 6.**
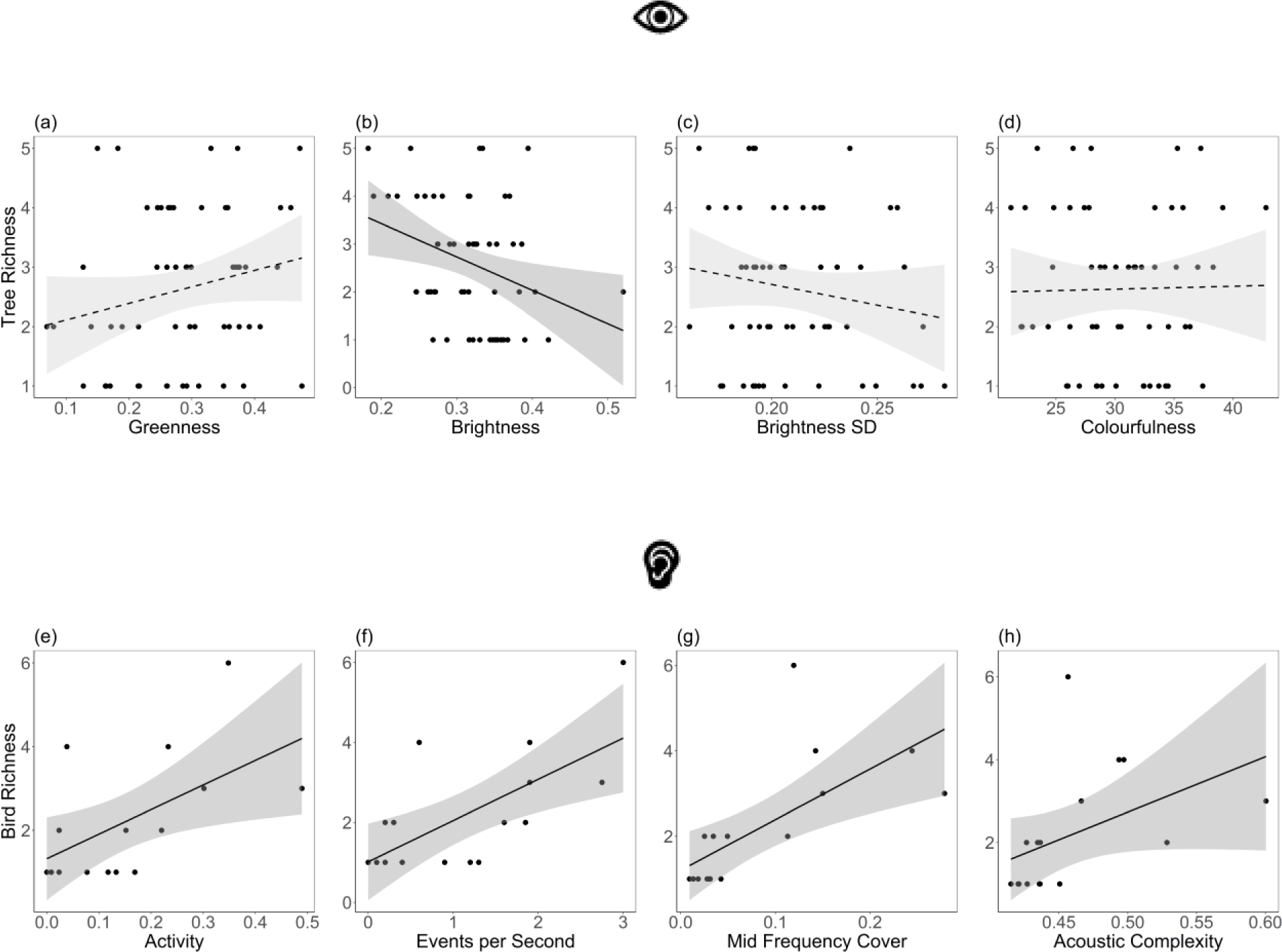
Relationships between the computed indices and actual diversity. Panels a-d show correlations for visual indices and tree richness, both derived from the photographs. Panels e-h show correlations for acoustic indices and bird richness, both derived from the audio recordings. Solid lines present significant correlations and dashed lines represent non-significant correlations, see details in Table 4. Shaded areas represent a 0.95 confidence interval.

## 4. Discussion

We could show in our study, focussing on the visual and acoustic sense, that perceived and measured forest biodiversity are significantly correlated - both for species richness and computed diversity indices. In particular, acoustic diversity can be well detected by people, while visual diversity may be harder to perceive. We identified perceived forest characteristics that could help bridge the gap between perceived and measured diversity: For the visual sense these were mainly structural parameters, e.g. vegetation density, light conditions, colours and forest structure. For the acoustic sense bird song characteristics, perceived physical properties such as volume, time of day as well as seasonal alterations and evoked emotions were the most frequently identified cues.

### 4.1 Perceived forest characteristics

Perception involves more than just the objectively measurable biophysical aspects. It also includes top-down processes like prior experiences, memories, knowledge, emotions, and individual physiological features that influence how we ultimately perceive things (e.g. Axelrod, 1974). We therefore identified subjective visual and acoustic forest characteristics in the open sorts to complement our understanding of what aspects, other than species richness, perceived diversity may consist of. The identified characteristics reflect both the bottom-up processing (e.g. colours of the forest, physical characteristics of the recording) and top-down processing (e.g. emotions, time of day). The significant environmental fitting analysis showed that the open sorts correlated with perceived visual and acoustic diversity (Fig. 1, Step 1), making the identified visual and acoustic forest characteristics likely candidates to better understand perceived visual and acoustic diversity.

For the visual sense, several of the perceived forest characteristics we identified partially align with work from Austen et al., (2021, 2023) where shared perspectives on forest diversity attributes were identified by asking people to sort illustration cards of four different taxa in different seasons. In our study, the most commonly used sorting criteria were vegetation density and light conditions, while in the Austen et al. (2021, 2023) studies these themes have been identified as meaningful concepts in sorts targeting trees in autumn only. Austin et al. (2021, 2023), however, presented single or few species on their illustration cards and no whole forest ecosystem as in the photographs of our study. As such, the participants’ attention in the Austen et al. (2021, 2023) studies may have been focused on characteristics of the depicted trees alone – rather than visual cues resulting from the interplay of several trees such as forest density or variations of light. As in our study, previous work also found that colour matters for perceived visual diversity (Austen et al., 2021, 2023; Hoyle et al., 2018). We further found that emotions and memories influenced the sorting strategies of participants which has also been reported by Austen et al. (2021, 2023) in their sorting experiment.

The perceived acoustic forest characteristics we identified support previous findings from studies that addressed bird sound perception (Ratcliffe, 2021; Ratcliffe et al., 2013, 2016, 2020). Participants in our study most often sorted the recordings according to bird song characteristics such as properties of the chirps, perceived melodic and repetitive patterns or complexity. With respect to restorative potential, qualitative (Ratcliffe et al., 2013, 2016) and quantitative (Ratcliffe et al., 2020) work identified similar acoustic properties as in the present study, i.e. physical characteristics such as the volume of bird sounds (Ratcliffe et al., 2013, 2020), people’s associations with the time of day or seasonal alterations (Ratcliffe et al., 2016), as well as expressed affective responses (Ratcliffe et al., 2013) or associated linkages with landscapes wherein bird sound might have been recorded (Ratcliffe et al., 2016). While we did not test the restorative potential of diversity in this study, our own work has previously shown that perceived acoustic diversity (Uebel et al 2021) and visual diversity (Rozario et al., 2024) are associated with restorative and mental well-being outcomes.

With our study, we provide evidence that there are measurable, bottom-up driven aspects of forests that people intuitively recognise when viewing forest photographs or hearing forest sound recordings, such as vegetation density or features of bird song, while we also identified top-down driven, subjective mental representations, such as evoked emotions. While top-down cognitive processes are hard to generalise, future research could elaborate on the link between the here identified biophysical forest features and how these translate into perceived biodiversity. For example, participants could be asked to sort photographs wherein species richness remains unchanged but other aspects, such as vegetation density, brightness, and colourfulness for perceived visual diversity, and melodic patterns or volume for perceived acoustic diversity are varied. Moreover, items that allow people to freely rate biodiversity according to their own understanding of biodiversity can be useful in studies where drivers of perceived diversity are tested in addition to their well-being effects. Rozario et al. (2024), for instance, asked participants how biodiverse they thought an environment was on a continuous scale from 0 (no biodiversity) to 100 (extremely biodiverse), and found that higher perceived diversity measured with this item was related to several mental well-being measures. At the same time, serving as an outcome variable, such items can be tested against potential influential factors of perceived diversity such as vegetation density for visual sense or volume for the acoustic sense. Future studies may further compare different psychological theories of perception, such as Schema Theory (Axelrod, 1974) or Gestalt theory (Koffka 1922), to better understand which of these theories best explain perceived biodiversity.

Importantly, our analysis revealed that for both the open visual and acoustic sorts, people also intuitively chose tree and bird richness as sorting criteria. This indicates that species richness - even if not in the first place - does stand out to people, rendering it an important aspect of perceived forest diversity.

### 4.2 Association between perceived and actual diversity

We found moderate to high associations between perceived diversity and species richness for both visual and acoustic stimuli (Fig.1, Step 2). This is in contrast to several studies reporting that participants cannot easily perceive biodiversity (Austen et al., 2021; Dallimer et al., 2012; Phillips & Lindquist, 2021; Rozario et al., 2024; Stobbe et al., 2022). The accuracy of people’s diversity estimations, however, might be influenced by whether they rate an environment’s diversity alone, or in comparison to other environments. Of the studies that identified discrepancies between perceived and actual diversity, two studies used cross-sectional measurements (Dallimer et al., 2012; Phillips et al., 2021), while in Rozario et al. (2024) participants were assigned to one environment only. In the present study, however, several forest environments were rated against each other, which enabled participants to directly compare diversity levels. Many studies reporting a good agreement between perceived and actual diversity also allowed participants to directly compare environments (e.g. Gao et al., 2019; Johansson et al., 2014; Simkin et al., 2020; Southon et al., 2018), thus supporting the assumption that the study design might be a crucial factor that determines the accuracy of diversity ratings, and our understanding of perceived diversity.

The participants’ ability to reflect bird richness in their ratings of acoustic diversity was stronger than their ability to reflect tree richness in their ratings of visual diversity. Compared to the photographs, the audio recordings were quite simple, as they did not contain any other noises when a bird vocalisation was not present. Therefore, the independent variable (bird species richness) was more easily identifiable by the participants. The forest images not only showed variations in tree richness but also understorey or forest structural attributes and were not otherwise against a blank background, as would be analogous for the acoustic recordings. We reasoned that the value of using stimuli from natural forest settings was more advantageous than less realistic stimuli. However, a replication of this study with simpler visual stimuli, through e.g. depicting tree species or close-up photos of their twigs against a neutral background (e.g. Austen et al., 2021, 2023; Hofmann et al. 2017), or conversely, more complex acoustic stimuli by combining bird vocalisations with sounds from insects, wind or water would be useful to confirm our conclusions. Furthermore, even experts struggle to visually differentiate certain tree species due to their similarity. Our photographs displayed combinations of species such as the European beach (*Fagus sylvatica*) alongside the common hornbeam (*Carpinus betulus*), which are hard to distinguish, particularly when presented on photographs and from afar. Future studies could therefore focus on highly conspicuous tree species alone or on the functional diversity of species traits (e.g., leaf traits such as evergreenness, or canopy traits such as branching patterns), which may be easier for people to detect. It could, however, also be that people’s general tendency to be more aware of animals compared to plants, also referred to as plant-blindness (Balding & Williams, 2016), explains our result pattern. That way, people may have been better in identifying bird richness as they allocated attentional capacities to birds opposed to participants who estimated visual diversity based on flora.

### 4.3 Diversity indices as proxies for perceived and actual diversity

We could identify computed diversity indices that can measure both perceived and actual diversity for the acoustic sense, but not for the visual sense. While we found a significant positive relationship between the greenness within photographs and their perceived visual diversity, greenness did not correlate with actual diversity, i.e tree richness. Similarly, the Brightness index was significantly negatively associated with actual diversity, but not related to perceived diversity (Fig.1, Step 3; Table 4).

The negative correlation between the Brightness index and actual diversity indicates that the photographs were darker in plots with higher tree species richness. This may be because forests with higher tree species richness can have greater canopy packing, and thus less light in the understory (Jucker et al., 2015).

The significant positive correlation between the visual index Greenness and perceived visual diversity, as well as the importance of vegetation density in the open sorts suggest that it is the amount of green vegetation that influences perceived visual forest diversity. Research by Dallimer et al. (2012) supports this interpretation: they reported a significant association between the tree cover of riparian areas and perceived species richness. Schebella et al. (2019) further investigated the linkages between biodiversity and mental well-being in urban green spaces, finding that several diversity metrics, including vegetation cover, were significantly associated with well-being. Interestingly, in their study, the well-being effects of naturalness, bird species richness, habitat diversity and structural heterogeneity all became insignificant when controlling for the effects of vegetation cover, thus underpinning the relevance of vegetation for human perception and human-biodiversity interactions more generally (Schebella et al., 2019).

No association was found between visual indices representing variations of light or colour and perceived diversity. Regarding the brightness indices, one explanation may be that despite being able to recognise variations of light, participants do not necessarily associate this with forest diversity. However, our index capturing light contrasts (Brightness SD) was also not significantly correlated with actual diversity, so the best explanation is likely that this index simply does not reflect either actual or perceived diversity. The Colourfulness index was also not associated with perceived diversity although it was frequently cited in the open sorts (Table 2). Our colourfulness values may not have successfully reflected perceived diversity ratings because of the time of year the photographs were taken. Our photos captured in late summer may lack the vibrant colours present in spring and autumn, influenced by flowering plants in the understory and changing foliage. Despite participants observing subtle variations in colour in the open sorts, the Colourfulness index would have reflected mostly green, and therefore likely did not vary sufficiently to significantly reflect variations in tree richness, or participants’ perceptions of forest diversity, in late summer. Although visual indices failed to reflect actual and perceived diversity in this study, attempts at different times of year would more thoroughly address the question of visual indices as proxies for actual and perceived diversity. A cross-seasonal comparison would, for instance, allow for specifically investigating the influence of visual indices within and across seasons. Following our results, it may be possible that greenness is the decisive perceived diversity attribute in summer via different shades of green and in winter through the differentiation between deciduous and evergreen species, while colour may more accurately explain perceived visual diversity in spring via flowers and blossoms and autumn via leaf colours and fruits. A larger study could then compare the influence of greenness and colourfulness for perceived diversity across seasons.

Regarding acoustic diversity indices, all four indices were significantly correlated with both perceived and actual diversity. This indicates that for the auditory sense, diversity indices are well-suited to reflect both people’s subjective perception of acoustic diversity and actual bird richness. Interestingly, the correlations between each acoustic index and perceived diversity ratings were descriptively higher than their correlations with actual bird species richness, suggesting that acoustic indices may be more effective in reflecting perceived diversity than actual diversity (although they succeed in significantly reflecting both in the present study). One reason for this may be that acoustic indices tend to level off when more than a handful of species are singing at the same time which was found for simulated (Gasc et al., 2015) and real soundscapes (Beason et al., 2023). Similarly, human perception of vocalising bird richness may be impeded, the more species sing together and the more birdsongs overlap. Future studies could specifically study soundscapes with high bird richness to see whether perceived diversity ratings reach a plateau when a certain number of birds vocalise together and test whether this pattern aligns with the one seen for acoustic indices.

### 4.4 Multisensory biodiversity experiences

Well-being effects obtained through experiencing a forest are of multisensory origin by nature. This fact is taken up in forest bathing – a mindfulness-based practice whereby people immerse themselves in forests using all five senses, including the visual and auditory sense but also haptics, smell and taste (Kotera et al., 2020). For instance, one would palpate the various structures of barks, smell the volatiles that different tree species emit or taste different berries or wild herbs. Only few studies so far have highlighted the importance of haptic (Lopez-Cotarelo Flemons et al., 2019), olfactory (Bentley et al., 2023; Hedblom et al., 2019; Schebella et al., 2020) and taste related nature experiences for well-being (Franco et al., 2017). The scarcity of scientific evidence for senses other than vision and hearing on biodiversity - health linkages necessitates more studies to assess how exactly biodiversity manifests through all five senses and their relevance for the nature-health nexus.

The present study strengthens our understanding of both perceived visual and acoustic biodiversity individually and how these align with actual diversity. Future studies might build upon our findings by testing combined audio-visual perception of biodiversity or even include all five senses and how perceived multisensory diversity mirrors and diverges from actual diversity. To our best knowledge, only one study investigated whether perceived diversity differs after exposure to uni-vs. multisensory diversity with the result that the unisensory experience of biodiversity resulted in higher perceived species richness (Schebella et al., 2020). Future studies should therefore elaborate on the complex interplay of the human senses in biodiversity experiences, i.e. whether the layering of senses would result in antagonistic effects as seen in Schebella et al. (2020) or additive or synergistic effects of sensory stimuli. Understanding how we perceive diversity may then help to design biodiverse healthscapes for wildlife and people alike.

### 4.3 Conclusion

Our study provides evidence that perceived diversity and actual species richness are correlated for the visual and acoustic sense. Especially with respect to acoustic signals, people can detect biodiversity well. Computed acoustic diversity indices provide a good proxy for actual and perceived bird species richness. However, the visual indices neither reflect actual nor perceived diversity well. We provide first insights into the nature of perceived visual and acoustic forest diversity, and we suggest conducting more systematic research to quantify the relationship of forest diversity attributes other than species richness with perceived diversity while also accommodating for the senses not covered in the present work.

Since perceived visual and acoustic diversity are linked to mental well-being, our findings have important implications for designing biodiverse, healthy environments or nature-based health interventions. We recommend the conservation and restoration of diverse forests, characterised by a variety of tree species and high structural diversity of vegetation. Such forests should include vertical layers that provide niches for different vocalising bird species. This approach serves as a nature-based solution that not only counteracts biodiversity loss but also enhances the forests’ potential to promote mental well-being, offering a cost-efficient measure for public health.

## Supporting information

Visual_and_Acoustic_Index_Computation

## Acknowledgements

We would like to express our deep gratitude to all participants who made this study possible. Thanks also to Dagmar Müller and Urte Roeber for their valuable input in conceptualising this study, to Emily Steinhart, Malin Wappelhorst and Oliver Stegmann for their help with collecting the data and to Emily Steinhart and Birte Peters for their help with analysing the data. Last but not least, we express our gratitude to the iDiv Data & Code Unit and particularly Ludmilla Figueiredo for assistance with curation and archiving of the dataset. This research was funded by the ERA-Net BiodivERsA project “Dr.FOREST” that investigates links between forest biodiversity and human health and wellbeing, with the national funders German Research Foundation (DFG-428795724, Germany), French National Research Agency (ANR, France), Research Foundation – Flanders (FWO, Belgium), Austrian Science Fund (FWF, Austria) and National Science Center (NCN, Poland, project no. 2019/31/Z/NZ8/04032), as part of the 2018-2019 BiodivERsA call for research proposals. KR, RRYO and AB gratefully acknowledge the support of iDiv funded by the German Research Foundation (DFG– FZT 118, 202548816). SM additionally receives financial support via the AkWamo project (2221NR050C): “Feasibility study - integration of (bio)acoustic methods for quantifying biological diversity in forest monitoring”, which is supported on the basis of a resolution of the German Bundestag with funds of the Federal Ministry of Food and Agriculture (BMEL) via the Agency of Renewable Resources (FNR) as project management agency of the BMEL for the funding program Renewable Resources. SMs contribution further benefited from the support of the Centre de Synthèse et d’Analyse sur la Biodiversité (CESAB) at the Fondation pour la Recherche sur la Biodiversité and the inspirational discussions among the Acoucene consortium.

## Author contributions

Conceptualisation and Design: Kevin Rozario, Taylor Shaw, Erich Schröger, Melissa Marselle, Rachel Rui Ying Oh, Aletta Bonn; Data collation: Kevin Rozario, Mateo Giraldo Botero; Formal analysis: Kevin Rozario, Taylor Shaw, Valentin Ștefan, Rachel Rui Ying Oh, Melissa Marselle, Mateo Giraldo Botero; Writing - first Draft: Kevin Rozario, Taylor Shaw; Writing - Review and Editing: All authors contributed critically to the drafts and gave final approval for publication.

## Data Availability Statement

All data will be made openly accessible at the institutional data repository of the German Centre for integrative Biodiversity Research (iDiv) as soon as it successfully underwent data curation. At the moment it is under review.

