## Appendix for "Perceived biodiversity: is what we measure also what we see and hear?"

### Appendix A1 Details on visual stimuli

We used standardised photos of natural forests, depicting different levels of tree species richness with 1-5 tree species (see Table A1 below). To focus on tree diversity uniformly across all different forests, we aimed to avoid photographing specimens showing blossoms, fruits, sprouting leaves, or signs of damage, disease or branch cuts (Hofmann et al., 2017). Similar to other studies, animals (Hofmann et al., 2017) and built elements were also not depicted in the photographs (Chiang et al., 2017; Grassini, Revonsuo, et al., 2019) to ensure the images were only depicting forest diversity. Regarding composition, or the sky/soil ratio and the arrangement of trees, we tried to follow suggestions by Svobodova et al. (2014, 2018) who found that subjects’ preference ratings of photos increase when having the horizontal line that bisects a scenery into sky and land within the upper or central part of a picture and placing central elements according to the Rule of Thirds. We thus placed the horizontal line around the centre or within the upper third of the picture, yet tried to avoid placing trees along either dividing line, as their presence may create subconscious preferences or draw disproportionate attention to species along that line.

**Table A1.**
Species (combinations) and number of photos per richness category for the visual sort.

| Richness Category | Species (compositions) | Number of photos |
| --- | --- | --- |
| 1 | *Acer pseudoplatanus (A)*  *Betula pendula (B)*  *Carpinus betulus (C)*  *Fagus sylvatica (F)*  *Picea abies (Pic)*  *Pinus sylvestris (Pin)*  *Quercus robur (Q)* | 2  1  2  4  1  2  1  _____  **13** |
| 2 | A+F  A+ *Fraxinus excelsior (Fr)*  A+Pic  B+C  C+Pin  C+Q  F+Fr  F+Q  Fr+Pic  Pic+Pin | 2  1  1  2  2  1  1  2  1  1  _____  **14** |
| 3 | A+F+Fr  A+F+Q  A+Fr+Pic  B+C+Q  B+C+Pic  B+C+Pin  C+Pic+Pin  C+Pic+Q  F+Fr+Pic  F+Pic+Pin | 2  1  1  1  3  1  1  1  2  1  _____  **14** |
| 4 | A+B+C+Pin  A+F+Fr+Pic  A+F+Fr+Q  A+F+Pic+Q  A+Fr+Pic+Q  B+F+Fr+Pic  B+Pic+Pin+Qro  C+Pic+Pin+Q | 1  1  2  1  1  2  1  2  _____  **11** |
| 5 | B+C+F+Pic+Pin  B+C+Pic+Pin+Q  B+F+Fr+Pic+Q | 1  2  2  _____  **5** |
|  | In total: | **57** |

### Appendix A2 Details on acoustic stimuli

We used acoustic recordings of natural bird assemblages in the same forest as for visual diversity with 1-6 birds singing (see Table A2 below). While acoustic forest diversity comprises more than just bird vocalisations, for example stridulating insects, croaking frogs or the sound of wind, we restricted the stimuli only to bird vocalisations because birds are highly vocal (and therefore commonly heard by people), and within the avian order, exhibit a wide diversity of vocalisations, even in a temperate climate. To standardise potential background sounds that can be observed in a forest (wind, rustling leaves, far-off animal sounds, anthropogenic noise), and to ensure that the sound stimuli were biologically relevant, we selected recordings of birdsong that were otherwise silent. We also varied characteristics within recordings in secondary ways, such as whether a vocalisation occurred early or late in the recording, and including complex vocalisations such as passerine birdsong, and simple vocalisations such as a crow’s ‘caw’. We did this to ensure we were not confounding participants’ ratings of acoustic diversity with a particular type of bird vocalisation, nor with the temporal sequence of silence and vocalisation.

**Table A2.**
Species (combinations) and number of recordings per richness category for the acoustic sort.

| Richness Category | Species (compositions) | complex/simple call type | Number of recordings |
| --- | --- | --- | --- |
| 1 | *Corvus corax (C)*  *Fringilla coelebs (F)*  *Fringilla coelebs (F)*  *Pyrrhula pyrrhula (P)*  *Regulus regulus (R) Turdus philomelos (TP)*  *Turdus viscivorus (TV)* | simple  simple  complex  simple  complex  simple  complex | 1  1  1  1  1  1  1  ________  **7** |
| 2 | *Erithacus rubecula (E)* + F  *Periparus ater (PA)* + unknown (u)  TV + u | complex  simple  simple | 1  2  1  ________  **4** |
| 3 | F + R + TP F + TP + *Troglodytes troglodytes (TT)* | complex  simple | 1  1  ________  **2** |
| 4 | E + F + PA + *Phylloscopus collybita* E + PA + TP + u | complex  complex | 1  1  ________  **2** |
| 6 | *Dryocopus martius* + E + F + *Sylvia atricapilla* + TT + u | complex | **1** |
|  |  | In total | **16** |

### Appendix A3 Summary of visual and acoustic index values.

| **Sensory pathway** | **Index** | **Description** | **Citation** | **Mean +- SD** | **Range** |
| --- | --- | --- | --- | --- | --- |
| Visual | Greenness | A quantification of the green values comprising a digital RGB photograph as one aspect of the Red-Green-Blue Vegetation Index. Higher values indicate more greenness. | Bendig et al., 2015; Lussem et al., 2018; Frey et al., 2019 | 0.287 ± 0.099 | 0.069 - 0.475 |
|  | Brightness | Converts image to grayscale, and quantifies how close to white the average pixel value is. Higher values indicate more brightness. | Frey et al., 2019; Menzel & Reese, 2021; Kardan et al., 2015; Ibarra et al., 2017 | 0.314 ± 0.061 | 0.183 - 0.520 |
|  | Brightness Standard Deviation | As a measure of light contrasts, this index takes the mean of the brightness index above and quantifies the amount of variation in light versus dark values present in the photograph. Higher values indicate stronger contrast. | Frey et al., 2019; Menzel & Reese, 2021; Kardan et al., 2015; Ibarra et al., 2017 | 0.211 ±  0.029 | 0.161 - 0.282 |
|  | Colourfulness | A quantification of colorfulness calibrated on the perception of participants based on the standard deviations and mean values within an opposing colour space defined along the red-green axis against the blue axis. Higher values indicate more colours. | Hasler & Süsstrunk (2003) | 30.403 ±  4.903 | 21.186 - 42.779 |
| Acoustic | Acoustic Complexity Index (ACI) | Designed to reflect complex sound, the ACI captures rapid variations in frequency and amplitude that are typical of biophony, i.e. biotic components of a soundscape (especially birdsong). This index typically does not respond to persistent sound such as machinery noise or buzzing insects. Higher values indicate greater biophony. | Pieretti et al., 2011; Towsey et al., 2017 | 0.459 ±  0.0495 | 0.415 - 0.600 |
|  | Events Per Second | Quantifies variations of quiet-to-loud events. The number of acoustic events per second in each noise-reduced frequency bin, where an event is counted each time the decibel value in a bin crosses the 3 dB threshold from lower to higher values, then divided by the number of seconds in the audio file. Higher values indicate more events per second in the audio recording. | Towsey et al., 2017 | 1.138 ± 0.956 | 0 - 3 |
|  | Activity | The fraction of values in the noise-reduced decibel envelope that exceed the threshold of 3 dB. Increases with more events generally in sound files. Higher values indicate more events in the audio recording. | Towsey et al., 2017 | 0.147 ± 0.141 | 0 - 0.489 |
|  | Mid Frequency Cover | The fraction of noise-reduced spectrogram cells that exceed 3 dB in the mid-frequency band (1000-8000 Hz), where biophony is typically present, including birdsong. Higher values indicate more biophonic events in the audio recording. | Towsey et al., 2017 | 0.083 ± 0.084 | 0.009 - 0.279 |

The R code used to compute these indices can be found in the supplementary .txt file (‘Visual_and_Acoustic_Index_Computation.txt’).

### Appendix A4 Correlations between diversity indices

**Table A41.**
Correlation coefficients of comparisons between visual diversity indices. All tests were Spearman correlation tests, unless denoted by a ⬨, indicating a Pearson correlation. Significant correlations are bolded and indicated as follows: < .05*, < .01**, <.001***.

|  | Greenness | Brightness | Brightness.SD | Colourfulness |
| --- | --- | --- | --- | --- |
| Greenness | 1 | **-0.45***** | 0.04 | **0.41**** ⬨ |
| Brightness | **-0.45***** | 1 | 0.11 | **0.36**** |
| BrightnessSD | 0.04 | 0.11 | 1 | 0.05 |
| Colourfulness | **0.41**** ⬨ | **0.36**** | 0.05 | 1 |

**Table A42.**
Correlation coefficients of comparisons between acoustic diversity indices. All tests were Spearman correlation tests. Significant correlations are bolded and indicated as follows: < .05*, < .01**, <.001***.

|  | Activity | EventsPerSecond | MidFreqCover | Acoustic Complexity |
| --- | --- | --- | --- | --- |
| Activity | 1 | **0.95***** | **0.76***** | **0.63**** |
| EventsPerSecond | **0.95***** | 1 | **0.74***** | **0.64**** |
| MidFreqCover | **0.76***** | **0.74***** | 1 | **0.89***** |
| Acoustic Complexity | **0.63**** | **0.64**** | **0.89***** | 1 |

### Appendix A5 First-order clusters of all five raters and their assignment to the second-order clusters

**Table A5.**
First-order clusters of all five raters and their assignment to the second-order clusters.

| 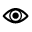 | | |
| --- | --- | --- |
| Rater | First-order cluster | Second order cluster |
| KR | density | density |
|  | light conditions | light |
|  | trunk | tree physical feature |
|  | colour | colour |
|  | forest floor vegetation | ground layer |
|  | diversity | diversity |
|  | structure | structure |
|  | perspective | perspective |
|  | mood | emotions |
|  | accessibility/transmissibility | ground layer |
|  | naturalness | structure |
|  | warmth | emotions |
| MM | density | density |
|  | colour | colour |
|  | order | structure |
|  | light | light |
|  | perspective | perspective |
|  | truck thickness | tree physical feature |
|  | inviting / welcoming | emotions |
|  | mood/feeling | emotions |
|  | accessibility | ground layer |
|  | height | structure |
|  | cultivated | structure |
|  | naturalness | structure |
|  | wellbeing | emotions |
|  | structure | structure |
|  | tree species | diversity |
| RRYO | density (tree/leaves) | density |
|  | ground layer | ground layer |
|  | knowledge (forest-related) | diversity |
|  | tree physical feature | tree physical feature |
|  | forest physical structure | structure |
|  | light | light |
|  | sensation | emotions |
| MGB | character or kind | OTHERS |
|  | colour | colour |
|  | density | density |
|  | effect or relationality | OTHERS |
|  | light conditions | light |
|  | measurement or dimension | measurement |
|  | surroundings | OTHERS |
|  | texture or materiality | OTHERS |
| OS | density | density |
|  | colour | colour |
|  | floor | ground layer |
|  | species | diversity |
|  | light | light |
|  | structure | structure |
|  | vegetation | structure |
|  | point of view | perspective |
|  | feels | emotions |
|  | cultivation | structure |
|  | diverse | diversity |
| 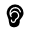 | | |
| rater | First-order cluster | Second order cluster |
| KR | bird interaction | bird interaction |
|  | landscape | landscape |
|  | bird song | bird song |
|  | birds' mood | vitality |
|  | time of day | time |
|  | salient sounds | salient sound |
|  | elements | OTHERS |
|  | intensity | vitality |
|  | bird abundance | bird abundance |
|  | bird species richness | bird species |
|  | mood | emotions |
|  | colour | memories / imagination |
|  | season | time |
|  | acoustic diversity | bird song |
|  | physical characteristics | physical characteristics |
|  | distance to sounds | physical characteristics |
|  | ratio bio-/geophony | salient sound |
|  | human company/fellowship | OTHERS |
|  | odour | memories / imagination |
|  | familiarity | memories / imagination |
|  | regularity | regularity |
| MM | river | landscape |
|  | chirping | bird song |
|  | liveliness | vitality |
|  | salient sounds | salient sound |
|  | sound intensity/volume | physical characteristics |
|  | amount of birds | bird abundance |
|  | bird species | bird species |
|  | bird species richness | bird species |
|  | mood / emotion | emotions |
|  | time (of day/season) | time |
|  | place | landscape |
|  | colour | memories / imagination |
|  | melody | bird song |
|  | complexity | regularity |
|  | background sounds | salient sound |
|  | smell | memories / imagination |
|  | tree density | landscape |
|  | structure / order | regularity |
| RRYO | Bird acoustics (Vitality, in the sense of distinguishing the living from the nonliving) | bird song |
|  | Landscape | landscape |
|  | Vitality | vitality |
|  | Sound characteristics (e.g. volume, timbre etc.) | physical characteristics |
|  | Sensation/mood | emotions |
|  | Time (of day or within a year) | time |
|  | Memories | memories / imagination |
| MGB | bird interaction | bird interaction |
|  | characteristics | physical characteristics |
|  | metric or sequence | physical characteristics |
|  | number of birds | bird abundance |
|  | sensation or feeling | emotions |
|  | sound singularity | bird song |
|  | surroundings or space experience | landscape |
|  | time of the day | time |
|  | time of the year | time |
| OS | activity | vitality |
|  | surrounding | landscape |
|  | bird song | bird song |
|  | time of day | time |
|  | others | OTHERS |
|  | volume | physical characteristics |
|  | abundance | bird abundance |
|  | species | bird species |
|  | species_abundance | Bird species / bird abundance |
|  | bird / rest | salient sound |
|  | emotion | emotions |
|  | season | time |
|  | melody | bird song |
|  | distance | physical characteristics |
|  | frequency | physical characteristics |

### Appendix A6 Environmental fitting analyses with actual diversity, perceived diversity and the diversity indices

Environmental fitting indicated that perceived visual similarity, i.e. the sorting pattern in the open visual sorts, was significantly associated with perceived visual diversity in the closed sorts (*p*<.001, *R^2^*=0.58; Fig.1, Step 1, Analysis 3), the Greenness Index (*p*<.001, *R^2^*=0.29) and the BrightnessSD index (*p*=0.02, *R^2^*=0.16). There was no significant relationship between perceived visual similarity and actual diversity (tree richness) and the visual indices Brightness and Colourfulness. Perceived acoustic similarity was significantly associated with perceived acoustic diversity (*p*<.001, *R^2^*=0.96), actual acoustic diversity (bird richness, *p*<.001, *R^2^*=0.85), as well as the indices Activity (*p*=0.04, *R^2^*=0.43), Events Per Second (*p*=0.01, *R^2^*=0.50), Mid Frequency Cover (*p*<.001, *R^2^*=0.82) and Acoustic Complexity (*p*=0.002, *R^2^*=0.62).

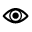

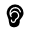

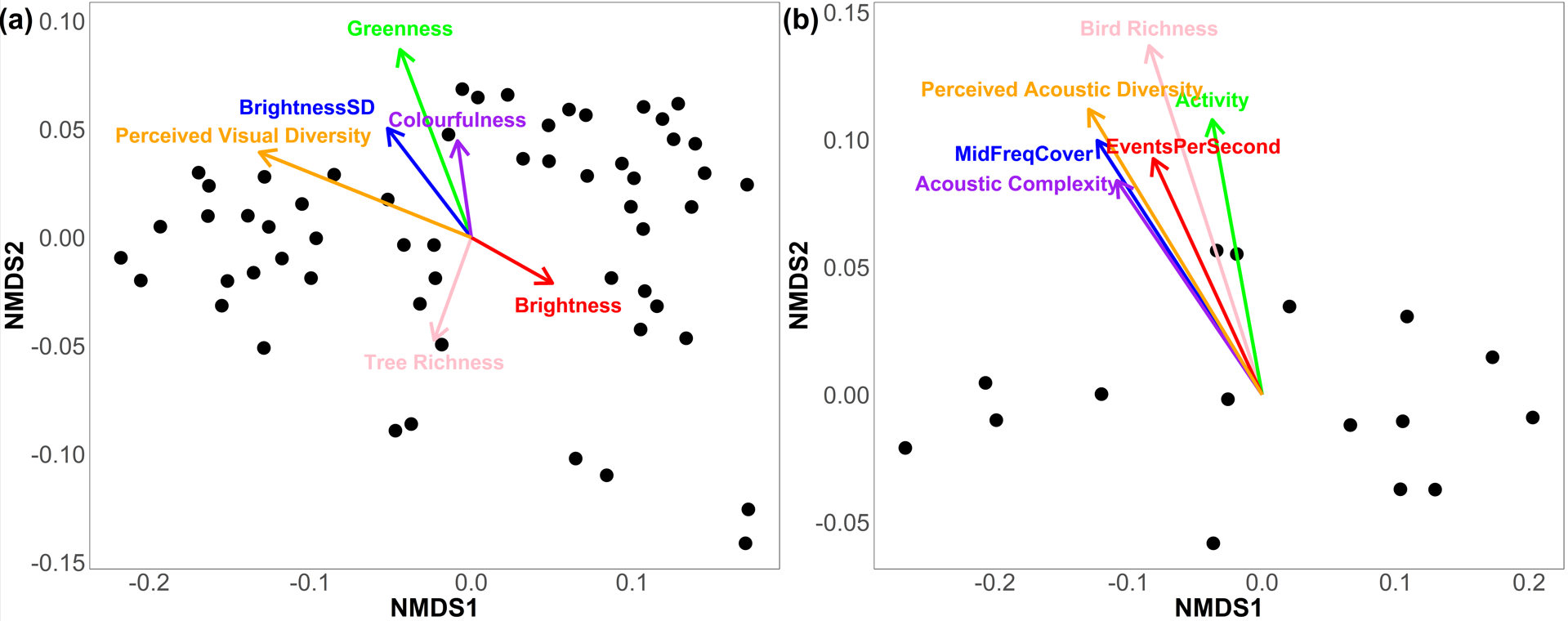

*Figure A6*. Non-metric multidimensional scaling (NMDS) plots resulting from the open visual (a) and open acoustic sort (b). Points represent photos and recordings, respectively. The closer two points (shorter distance), the more similar photos and recordings were perceived as. Environmental fitting was conducted to see whether perceived diversity, actual diversity and the diversity indices align with the produced NMDS solutions (stress values: visual: 0.06; acoustic: 0.04). The arrows illustrate the direction and strength of the associations between perceived diversity, actual diversity, the diversity indices and the NMDS solutions.

### Appendix A7 References Appendix

Chiang, Y. C., Li, D., & Jane, H. A. (2017). Wild or tended nature? The effects of landscape
location and vegetation density on physiological and psychological responses.
*Landscape and Urban Planning*, *167*(June), 72–83.
<https://doi.org/10.1016/j.landurbplan.2017.06.001>

Grassini, S., Revonsuo, A., Castellotti, S., Petrizzo, I., Benedetti, V., & Koivisto, M. (2019).
Processing of natural scenery is associated with lower attentional and cognitive load
compared with urban ones. *Journal of Environmental Psychology*, *62*(January), 1–11.
<https://doi.org/10.1016/j.jenvp.2019.01.007>

Svobodova, K., Sklenicka, P., Molnarova, K., & Vojar, J. (2014). Does the composition of
landscape photographs affect visual preferences? The rule of the Golden Section and
the position of the horizon. *Journal of Environmental Psychology*, *38*, 143-152.

Svobodova, K., Vojar, J., Sklenicka, P., & Filova, L. (2018). Presentation matters: Causes of
differences in preferences for agricultural landscapes displayed via photographs and
videos. *Space and Culture*, *21*(3), 259-273.
