## Supplementary material for "Perceived biodiversity: is what we measure also what we see and hear?": Visual_and_Acoustic_Index_Computation

#Visual index computation

## Packages

library(Rcpp)
library(terra)

## Directory with image files
base_dir <- #[load filepath here for folder with images]
  
  ## Functions
  greenness <- function(img){ # actually the RGBVI from https://www.int-arch-photogramm-remote-sens-spatial-inf-sci.net/XLII-3/1215/2018/isprs-archives-XLII-3-1215-2018.pdf
    rgbvi <- ((img[[2]]*img[[2]])-(img[[1]]*img[[3]]))/((img[[2]]*img[[2]])+(img[[1]]*img[[3]]))
    return(mean(values(rgbvi), na.rm = T))
  }

brightness <- function(img) { # calculates the image brighness and brighness sd
  grey <- mean(img/255)
  return(c(mean = mean(values(grey)),sd = sd(values(grey))))
}


##colourfulness

# David Hasler, Sabine E. Suesstrunk, "Measuring colorfulness in natural
# images," Proc. SPIE 5007, Human Vision and Electronic Imaging VIII, (17
# June 2003); doi: 10.1117/12.477378
# Efficient colorfulness metric
colorfulness <- function(img){
  rg <- img[[1]] - img[[2]]
  yb <- 0.5*(img[[1]] + img[[2]]) - img[[3]]
  
  sig <- max(c(mean(sqrt(apply(as.array(rg),1,sd)^2 + apply(as.array(yb),1,sd)^2)),
               mean(sqrt(apply(as.array(rg),2,sd)^2 + apply(as.array(yb),2,sd)^2))))
  mue <- max(c(mean(sqrt(apply(as.array(rg),1,mean)^2 + apply(as.array(yb),1,mean)^2)),
               mean(sqrt(apply(as.array(rg),2,mean)^2 + apply(as.array(yb),2,mean)^2))))
  
  cfs <- sig + 0.3 * mue
  return(cfs)
}


## Process images
# create a loop over images in base_dir and write results in the same folder
setwd(base_dir)
jpgs <- list.files(pattern = '*.JPG')

results <- data.frame(file = jpgs, greenness = NA, brightness = NA, brightness.sd = NA,
                      colorfulness = NA)
i <- 1
for(j in jpgs){
  img <- suppressWarnings(rast(j))
  results[i,2:ncol(results)] <- c(greenness(img), brightness(img), colorfulness(img))
  i <- i+1
}

write.csv(results,'visual_indices.csv', row.names = F)


###############################################

#Acoustic index computation

rm(list=ls())
require(stringr) #Simple, Consistent Wrappers for Common String Operations
require(chron)
require(soundecology)
require(tuneR)

#####FIRST THE TOWSEY ANALYSIS PACKAGE NEEDS TO BE DOWNLOADED FOR THIS SCRIPT TO CALL UPON IT
#URL: https://ap.qut.ecoacoustics.info/


# set working directory

setwd("F:/sorting_task/rawfiles")

folders_all <- list.dirs(path="F:/sorting_task/rawfiles")   #list of folder names
j=5
path=NULL

for(k in 1:length(folders_all)){
  if(k==j)
  {
    break;
  } else
  {
    path<-paste(folders_all[k], sep="")
    setwd(path)
    # Get a list of audio files inside the directory
    # (Get-ChildItem is just like ls, or dir)
    files_all2 <- list.files(path=path, pattern = "*.wav", recursive=FALSE)
    
    # The directory to store the results
    
    base_output_directory <-paste() #create a towsey_output folder for results to be stored
    
    # iterate through each file
    for(file in files_all2) {
      message("Processing ", file)
      # get just the name of the file
      file_name <- basename(file)
      # make a folder for results
      output_directory <- normalizePath(file.path(base_output_directory, file_name))
      dir.create(output_directory, recursive = TRUE)
      # prepare command
      command <- sprintf('audio2csv "%s" "C:/AP/ConfigFiles/Towsey.Acoustic.48khz_10sec_DrFOREST_sort_task.yml" "%s" ', file, output_directory) #NOTE: this .yml file was edited to handle recordings of 10sec lengths and up to 24kHz. Adjust parameters and save new .yml file (with new name), according to your recording length and freq range
      # finally, execute the command
      system2('C:/AP/AnalysisPrograms.exe', command)
    }
  }
}


#Indices are calculated. Code below allows you to compile them.

setwd() #set working directory to your towsey_output folder
folders <- list.files(pattern = 'Towsey.Acoustic.Indices.csv', recursive = TRUE)
blank <- data.frame()

for (i in 1:length(folders)){
  path <- paste('[##insert your filepath for towsey output folder##]', folders[i], sep = '')
  file <- read.csv(path)
  blank <- rbind(blank, file)
}
final <- blank

final$filename2 <- paste(final$FileName, '.wav', sep = '')
write.csv(final, 'acoustic_ais.csv')
